# The epigenetic reader MORC3 is required for T cell development in the thymus

**DOI:** 10.1101/2025.03.05.641591

**Authors:** Veronica Della Chiara, Maja Vukic, Jihed Chouaref, Laura Garcia-Perez, Gita Naber, Kelly K. D. Vonk, Christ Leemans, Artemiy Kovynev, Zhenhui Zhong, Cor Breukel, Sandra G. Vloemans, Kirsten Canté-Barrett, Karin Pike-Overzet, Steven E. Jacobsen, Frank J. T. Staal, Lucia Daxinger

## Abstract

**Cell fate decisions are orchestrated by an interplay of epigenetic factors in collaboration with transcriptional activators and repressors. The epigenetic reader MORC3 is known as a silencer of endogenous retroviruses in mouse embryonic stem cells. Here, we show that mouse MORC3 is expressed in the thymus and reveal that MORC3 loss of function leads to a severe arrest in T cell development at the DN1 stage, accompanied by an expansion of natural killer and myeloid cells. MORC3 function in thymus depends on its ATPase and its ability to bind H3K4me via the CW-type zinc finger domain. We find altered chromatin accessibility at regulatory elements of key T cell transcription factors in DN1 cells and show that re-expressing TCF1 in MORC3-deficient progenitor cells, rescues T cell development. Our work indicates a novel function for MORC3 in modulating regulatory sites essential for T cell lineage commitment at the level of chromatin.**

## Introduction

Cell fate decisions during development are not only directed by changes in gene expression controlled by transcription factors and repressors, but also by epigenetic modifications, such as DNA methylation and histone modifications. Such fate decisions can be studied in great detail for development of T lymphocytes, because genotype-phenotype relationships are relatively well understood^1,2^. During T cell development, hematopoietic progenitor cells migrate from the bone marrow to the thymus. Once in the thymus, these cells go through a series of differentiation stages, ultimately leading to the generation of mature and diverse T-cell populations that can be distinguished based on a large number of well characterized surface molecules^3^.

Progenitor cells that enter the thymus have the potential to develop into many different immune cell types, not only T cells. The irreversible potential to only form T lineage cells is termed commitment and involves a number of signaling routes and transcription factors. While Notch signaling and transcription factors such as TCF1^4^ and BCL11B^5^ are well characterized as drivers of T cell lineage commitment^6,7^, the epigenetic regulation of this process remains poorly understood^8^. However, the interplay between epigenetic factors and transcriptional regulators ensures the appropriate activation and repression of genes required for T cell lineage commitment. Hence it is important to get better insights into the epigenetic regulators that play a specific role in T cell development.

Histone marks are post-translational modifications predominantly found at the amino-terminal tails of histone proteins and include acetylation, methylation, phosphorylation and ubiquitination. The enzymes responsible for the deposition and removal of these marks are referred to as writers and erasers, respectively. In addition, epigenetic readers recognize specific modifications to mediate downstream effects. One of these readers is MORC3, a member of the highly conserved Microrchidia (MORC) family of nuclear proteins that comprises four members, MORC1-4, in mammals^9,10^. MORC proteins contain a conserved GHKL-type ATPase domain at their N-terminus, a PHD-X/ZF-CW domain and C-terminal coiled-coil domains that can include sites for SUMOylation. Mammalian MORC proteins show tissue specific expression patterns, which are important for development and have roles in DNA repair pathways and transposable element silencing^11,12^. Human MORCs have been linked to developmental and autoimmune diseases as well as cancer^13,14^, emphasizing their critical cellular functions.

In mice, *Morc3* knockout results in perinatal lethality^15,16^, although the cause of death is unknown. We previously identified MORC3 from a forward genetic screen for epigenetic regulators in the mouse^16^, and in mouse embryonic stem cells (mESCs) and male germ cells, MORC3 is required for silencing of transposable elements^16-18^. *Morc3* expression is high in blood and immune cells (https://www.ebi.ac.uk/gxa/genes/) and recently, MORC3 has been identified as a negative regulator of the interferon pathway in a human monocytic cell line^19^, suggesting that it is functionally important in immune cells. Structural and biochemical studies have shown that MORC3 is an epigenetic reader of H3K4me via its CW-domain^20,21^ and MORC3 can localize to two distinct chromatin states, the active histone mark H3K4me3 and the repressive modification H3K9me3^16,17,21^. Knockout of *Morc3* in mESCs leads to a gain in chromatin accessibility at H3K9me3 decorated endogenous retroviruses and upregulation of these elements. However, MORC3s function at active chromatin has so far remained elusive.

Here, we show that MORC3 loss of function mice have a small thymus due to a hematopoietic defect in T cell development at the stage when irreversible lineage commitment occurs. This effect is dependent on MORC3 targeting to chromatin via its ATPase and CW zinc finger domains, and is associated with high levels of apoptosis. While Notch signaling is largely unaffected, we found reduced chromatin accessibility of enhancers for T cell commitment factors such as TCF1 and BCL11B, while regulatory elements of non-T cell lineage factors show increased accessibility in MORC3 deficient thymus. Indeed, forced expression of TCF1 or BCL11B can rescue the developmental arrest caused by MORC3 loss of function. We propose that MORC3, through its ability to bind to active chromatin, helps to establish and maintain the unique gene expression patterns that define the identity and differentiation of T cells by regulating chromatin accessibility of T cell transcription factor regulatory elements.

## Results

### *Morc3^MD41/MD41^* mice exhibit abnormal T cell development

Using our previously reported *Morc3^MD41^* mouse model carrying a MORC3 nonsense mutation that results in a premature stop codon at Tyr327 in the ATPase domain (Fig. 1A), we set out to investigate the potential role of MORC3 in the immune system. Considering that *Morc3^MD41^* homozygotes are perinatally lethal, we focussed on the embryo. We found that MORC3 is expressed in embryonic immune organs including fetal liver and spleen and is particularly abundant in the thymus (Fig. 1B). As expected for a nonsense mutation, MORC3 protein levels were virtually absent in *Morc3^MD41^* homozygotes when measured by western blot in embryonic thymus (Fig. 1C), consistent with a loss of function. At a macroscopic level, while fetal liver and spleen showed no differences between genotypes, we noticed a much smaller thymus in *Morc3^MD41^* homozygous mutant embryos (Fig. 1D and Supplementary FigS1A). This was reflected in an almost 100-fold lower thymocyte cell count in *Morc3^MD41^* homozygotes, when compared to wild type (WT) and heterozygous littermates (Fig. 1D). These phenotypic features of *Morc3^MD41^*homozygous thymi prompted us to characterize thymocyte development in detail.

**Figure 1.**
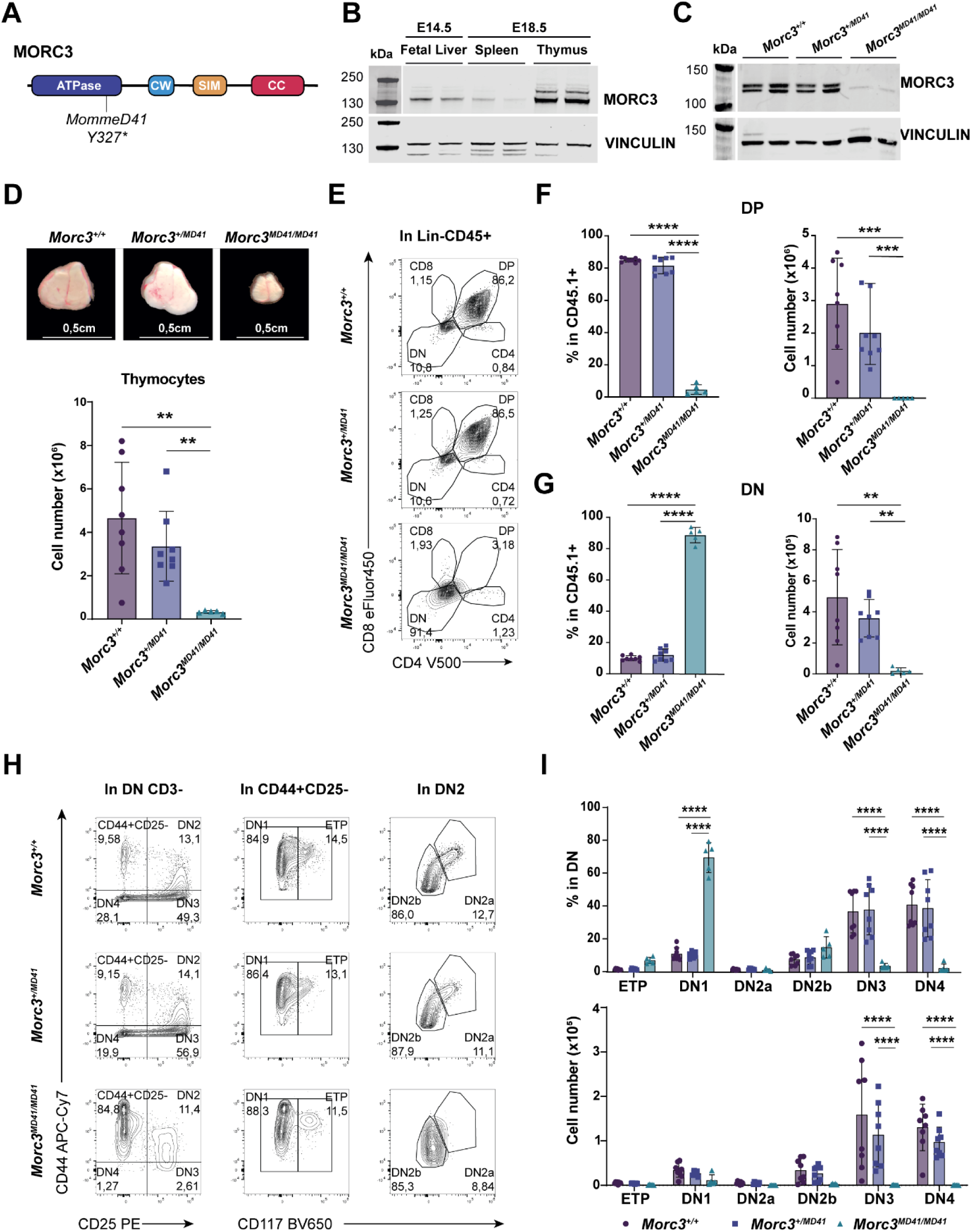
MORC3 deficiency leads to impaired T cell development. (A) Schematic representation of MORC3 protein, its different domains and the *MommeD41* mutation. (B) Western blot showing MORC3 protein levels in E14.5 fetal liver and E18.5 spleen and thymus, two *Morc3^+/+^* biological replicates. VINCULIN was used as a loading control. (C) Western blot showing MORC3 protein levels in E18.5 thymus, two biological replicates per genotype. VINCULIN was used as a loading control. (D) (top) Representative macroscopic pictures of thymus from E18.5 *Morc3^+/+^, Morc3^+/MD41^* and *Morc3^MD41/MD41^* littermates (scale bars, 0,5cm). (bottom) Total cell count of E18.5 thymus from *Morc3^+/+^, Morc3^+/MD41^* and *Morc3^MD41/MD41^* littermates (n = 8 *Morc3^+/+^*, n = 8 *Morc3^+/MD41^*, n = 5 *Morc3^MD41/MD41^*). Data are presented as mean with dot, square or triangle as individual values; error bar represents standard deviation (SD). (E) Representative flowcytometry plots of T cell subsets in *Morc3^+/+^, Morc3^+/MD41^* and *Morc3^MD41/MD41^* E18.5 thymus. CD4 and CD8 were used as markers to define DN, DP and single positive CD4+ and CD8+. (F) (left) Bar plot depicting the percentage of CD4+CD8+ Double Positive (DP) within CD45.1+ cells (pre-gated on SSC-A/FSC-A, single, live, Lin-) and (right) total cell number in E18.5 thymus (n = 8 *Morc3^+/+^, n* = 8 *Morc3^+/MD41^*, *n* = 5 *Morc3^MD41/MD41^*, 3 independent experiments). Data are presented as mean with dot, square or triangle as individual values; error bar represents standard deviation (SD). (G) (left) Bar plot showing the percentage of CD4-CD8-Double Negative (DN) within CD45.1+ cells (pre-gated on live, Lin-, CD3-) and (right) total cell number in E18.5 thymus (n = 8 *Morc3^+/+^, n* = 8 *Morc3^+/MD41^*, *n* = 5 *Morc3^MD41/MD41^*, 3 independent experiments). Data are presented as mean with dot, square or triangle as individual values; error bar represents standard deviation (SD). For panels D, F and G statistical significance was determined using two-sided *t* test. (H) Representative flowcytometry plots of DN cells in *Morc3^+/+^, Morc3^+/MD41^* and *Morc3^MD41/MD41^* E18.5 thymus. In the left column, CD25 and CD44 were used as markers to define DN1, DN2, DN3, DN4 T cell subsets within CD4-CD8-CD3-cells. In the middle column, c-kit was used as marker to define ETP and DN1 within CD44+CD25-cells. In the right column, c-kit was used as marker to define DN2a and DN2b within DN2 cells. (I) (top) Bar plot showing the percentage of DN cells in CD45.1+ cells (pre-gated on SSC-A/FSC-A, single, live, Lin-) and (bottom) total cell number in E18.5 thymus. (n = 8 *Morc3^+/+^, n* = 8 *Morc3^+/MD41^*, *n* = 5 *Morc3^MD41/MD41^*, 3 independent experiments). Data are presented as mean with dot, square or triangle as individual values; error bar represents standard deviation (SD). Statistical significance was determined using two-way ANOVA for multiple comparisons. For all figures: * P ≤ 0.05, ** P ≤ 0.01, *** P ≤ 0.001, **** P ≤ 0.0001.

Flowcytometry analysis of E18.5 *Morc3^MD41^* homozygous thymocytes revealed an almost complete absence of the major subpopulations of thymocytes, the Double Positive (CD4+CD8+, DP) cells, in both percentage and cell number (Fig. 1E-F). The drastic decrease of DP thymocytes was accompanied by a 9-fold increase in percentage of Double Negative (DN) (CD4-CD8-) cells (∼ 70% of total leukocytes in *Morc3^MD41/MD41^*, ∼15% in WT and *Morc3^+/MD41^*) in *Morc3^MD41/MD41^* thymus, indicating a developmental arrest in the DN subset (Fig. 1G). This increase in percentages was not seen in absolute cell numbers, because of the low overall thymocyte numbers observed in *Morc3^MD41^* homozygotes, when compared to WT and heterozygotes (Fig. 1G). We next performed a detailed immunophenotyping of DN subsets to gain insight into the impact of MORC3 loss on early T cell development (Fig. 1H). Within the DN compartment, a significant increase in the percentage of DN1 (CD44+CD25-ckit-) cells was observed in *Morc3^MD41/MD41^* thymi, pinpointing the developmental arrest to this most immature thymocyte stage. Notably, DN2a (CD44+CD25+ckit+) and DN2b (CD44+CD25+ckit-) cell counts and proportions in *Morc3^MD41/MD41^* embryos were comparable with WT and heterozygous counterparts, highlighting that a few early thymocytes were able to bypass this first major DN1 block in *Morc3^MD41^* homozygous mutants. Nevertheless, *Morc3^MD41/MD41^* thymocytes exhibited a severe depletion of DN3 (CD44-CD25+) and DN4 (CD44-CD25-) cell populations, evident in both their relative proportions and absolute cell counts, suggesting an additional developmental arrest at the DN3 stage (Fig. 1I). Collectively, these data show that MORC3 plays a crucial role in T cell development, particularly at the transition from DN1 to DN2 and culminating in a complete block at the DN3 stage.

To better understand the reason for the low thymocyte numbers in *Morc3^MD41^*homozygotes, we measured apoptosis and cell cycle in the various stages of early T cell development. While some level of apoptosis is intrinsic to T cell development due to selection processes, we found that the percentages of cells undergoing apoptosis were significantly higher in *Morc3^MD41^* homozygous thymocytes, increasing to ∼60% in DN2 cells (Fig. 2A and Supplementary FigS2A). This high level of apoptosis was accompanied by increased proliferation rates in all thymocyte populations analysed (Fig. 2B and Supplementary FigS2B), probably as a compensatory mechanism, but this did not lead to correction to normal cell numbers. These data show that MORC3 is required for early thymocyte survival.

**Figure 2.**
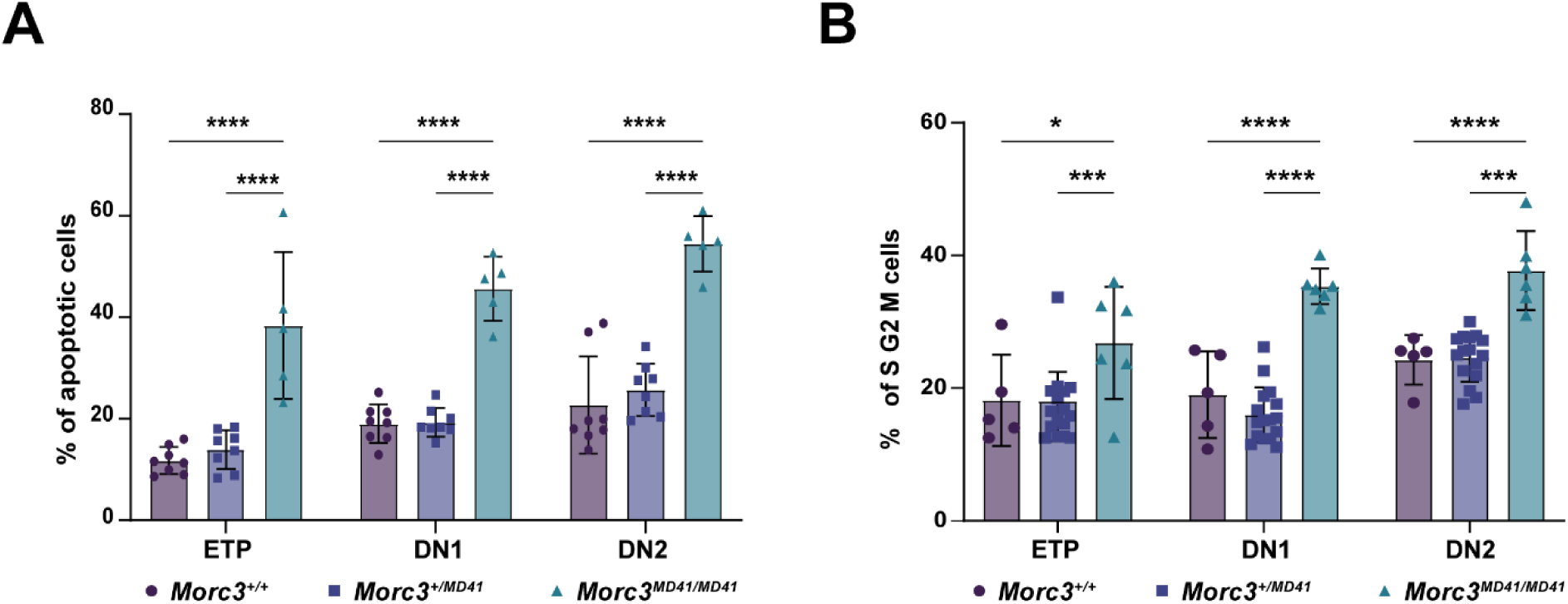
MORC3 is critical for T cell survival and proliferation. (A) Percentage of apoptotic cells (7AAD-, AnnexinV+) within ETP, DN1 and DN2 T cell subpopulations (n = 8 *Morc3^+/+^*, n = 8 *Morc3^+/MD41^*, n = 5 *Morc3^MD41/MD41^*, 3 independent experiments). Data are presented as mean with dot, square or triangle as individual values; error bar represents standard deviation (SD). (B) Percentage of cells in S, G2 and M phase of cell cycle (Ki67+, 7AAD+) within ETP, DN1 and DN2 T cell subpopulations (n = 5 *Morc3^+/+^, n* = 16 *Morc3^+/MD41^*, *n* = 6 *Morc3^MD41/MD41^*, 3 independent experiments). Only populations with ≥200 cells were included in the analysis. Data are presented as mean with dot, square or triangle as individual values; error bar represents standard deviation (SD). For all panels: statistical significance was determined using two-way ANOVA for multiple comparisons. P≤0.05, **P≤0.01, ***P≤0.001, ****P ≤ 0.0001.

### *Morc3^MD41/MD41^* thymus shows an expansion of myeloid cells and NK cells but not B cells

To gain insight into the molecular mechanisms underlying the developmental arrest in *Morc3^MD41/MD41^*thymus, we employed single cell RNA-sequencing (scRNA-seq) to capture cellular heterogeneity and gene expression dynamics in the thymus (Fig. 3A). Two samples (one *Morc3^MD41/+^* and one *Morc3^MD41/MD41^*; *Morc3^+/+^* and *Morc3^MD41/+^* tymi are morphologically indistinguishable on the day of dissection and processing for scRNA-seq Fig. 1D) were sequenced and cell types were identified based on the expression of known marker genes derived from previous publications^22,23^ (Supplementary FigS3A-B). UMAP visualization of the scRNA-seq data underscored the severe arrest at the DN1-2 stages of thymocyte development, with an absence of DP and a paucity of DN3 cells in *Morc3^MD41^* homozygous compared to heterozygous thymus (Fig. 3B and Supplementary FigS3C-D). We also observed an expansion of myeloid and natural killer (NK) cells, which are normally minor subpopulations in the thymus but now substantially increased in cell numbers and proportions in the homozygotes (Fig. 3B and Supplementary FigS3C-E), due to the absence of more mature thymocytes. Flowcytometry analysis of immune cell lineages confirmed the abnormal cellular distribution in independent samples. We found increased percentages of myeloid and NK cells in *Morc3^MD41^* homozygous thymi, while we did not observe any differences in the proportion of these subpopulations between WT and *Morc3^MD41^* heterozygotes (Fig. 3C and Supplementary FigS4). This is somewhat reminiscent of the phenotype observed in mice with targeted mutations in *Notch1^24^* or important transcription factors (TFs) that help establish the T cell gene programme, such as TCF1. However, in these mutants there is also an accumulation of B cells in the thymus^24^, which we did not observe (Fig. 3B-C), suggesting that Notch signalling proceeds relatively normal in absence of MORC3.

**Figure 3.**
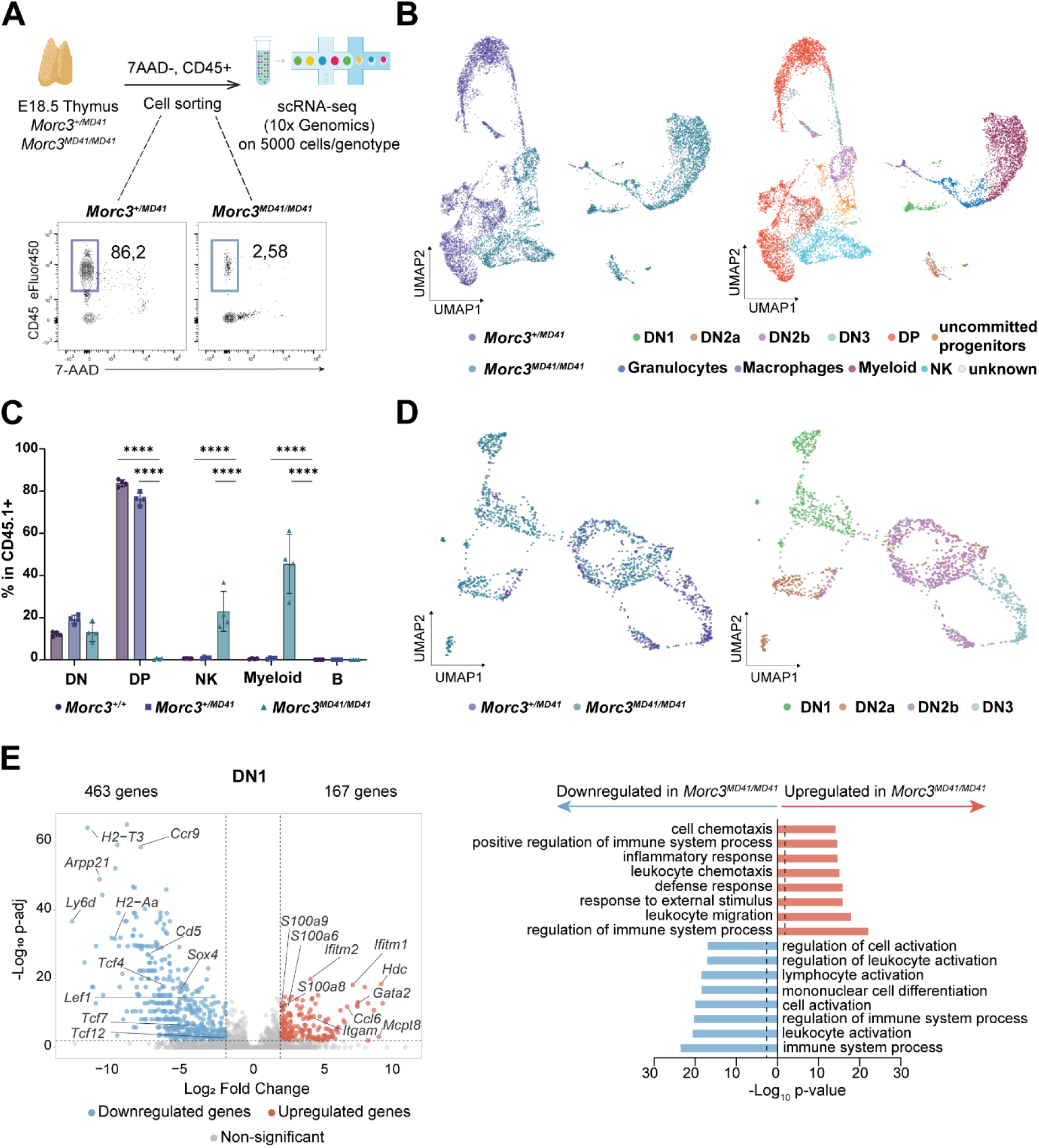
Aberrant accumulation of myeloid and natural killer (NK) cells and dysregulated transcriptome in *Morc3^MD41/MD41^* thymus. (A) Schematic representation of scRNA-seq experimental set-up and flow cytometry plots of 7AAD-CD45+ *Morc3^+/MD41^*and *Morc3^MD41/MD41^* thymocytes that were sorted and sequenced. (B) UMAP representations showing integrated live leukocytes in *Morc3^+/MD41^* and *Morc3^MD41/MD41^*E18.5 thymus where cells are colored by (left) genotype and (right) assigned cell type. Each dot represents a single cell. (C) Bar plots indicating the percentage of DN (Cd11b-Gr1-NK1.1-Cd11c-CD3-CD4-CD8-), DP (Cd11b-Gr1-NK1.1-Cd11c-CD3+CD4+CD8+), NK (Cd11b-Gr1-NK1.1+Cd11c-CD3-CD4-CD8-), myeloid (Cd11b+Gr1+NK1.1-Cd11c-CD3-CD4-CD8-) and B cells (Cd11b-Gr1-NK1.1-Cd11c-CD3-CD4-CD8-B220+CD19+) within CD45.1 cells in *Morc3^+/+^, Morc3^+/MD41^* and *Morc3^MD41/MD41^* E18.5 thymus (n = 4 biological replicates/genotype, 2 independent experiments). Data are presented as mean with dot, square or triangle as individual values; error bar represents standard deviation (SD). All cells are pre-gated on SSC-A/FSC-A, singles, live cells (7AAD-), CD45.1+. Statistical significance was determined using two-way ANOVA for multiple comparisons, *p≤0.05, **p≤0.01, ***p≤0.001, ****p≤0.0001. (D) UMAP representations showing integrated DN cell subsets in *Morc3^+/MD41^* and *Morc3^MD41/MD41^* after subset analysis and cells are colored by (left) genotype and (right) assigned cell type. Each dot represents a single cell. (E) (left) Volcano plot showing differentially expressed genes between *Morc3^+/MD41^* and *Morc3^MD41/MD41^* DN1 cells. Genes with |log2 fold change|> 2 and p-adj < 0.01 were considered significant. Red dots indicate upregulated genes in *Morc3^MD41/MD41^* DN1 cells. Blue dots indicate downregulated genes in *Morc3^MD41/MD41^* DN1 cells. Values on top of the plot denote number of upregulated and downregulated genes. The y-axis shows -log10 p-adj and the x-axis log2 fold change. Horizontal dashed line indicates -log p-value of 0,01 (=2), vertical dashed lines indicate log fold change of ±2. (right) Bar graph depicting Gene Ontology enrichment of biological processes in significantly differential gene sets in the DN1 cell cluster. Dashed lines indicate the -log p-value of 0,01.

Next, to identify differentially expressed genes and molecular pathways linked to the *Morc3^MD41/MD41^* early T cell developmental block we performed pseudo-bulk gene expression and Gene Ontology (GO) analyses on sub-setted (DN1-DN3) thymocyte populations (Fig. 3D-E and Supplementary FigS5A-D). We observed that expression of known Notch target genes (*Hes1*, *Dtx1, Ptcra, Il2ra, Id2, Nrarp, Myc, Notch1,2,3)* is largely unaffected in *Morc3^MD41^* homozygotes (Supplementary File 1). On the other hand, we found downregulation of lymphoid specific genes, including the key T cell genes *Tcf7*, *Bcl11b*, *Lck*, *Gata3* and *Rag1/2*, and upregulation of genes involved in inflammatory (*Ifitm1, Ifitm2, Il13*), migration processes (*S100a8, S1009a, Ccl6, Cxcr6*) and genes associated with myeloid cells (*Spi1, Gata2, Cpa3, Mcpt, Cma1*) in *Morc3^MD41^* homozygous thymus in all (DN1-DN3) sub-setted populations (Fig. 3E, Supplementary FigS5C-D and Supplementary File 1). Consistently, we found enrichment of genes linked to inflammatory response and lymphocyte development among the up- or downregulated genes, respectively (Fig. 3E and Supplementary FigS5C-D). Transcriptional dysregulation was present in *Morc3^MD41/MD41^* DN1 cells (Fig. 3E and Supplementary File 1), indicating that a deviation from the typical transcriptional program of T cells occurs even in the initial phases of thymocyte development. A bulk RNA-seq experiment carried out in WT and *Morc3^MD41/MD41^* embryonic thymus supported these observations. We found 3208 significantly differentially expressed genes (Supplementary FigS6A and Supplementary File 1) with numerous T cell marker genes among the 1446 downregulated, and myeloid genes among the 1762 upregulated genes in *Morc3^MD41/MD41^* thymus (Supplementary FigS6B and Supplementary File 1), and including *S100a8/a9*, *Rag1/2* and *Tcf7* (Supplementary FigS6C). Combined, these data show that loss of MORC3 function leads to transcriptional dysregulation of key immune cell TFs and their target genes and accumulation of myeloid cells in embryonic thymus.

### The *Morc3^MD41/MD41^* T cell defect is cell intrinsic

Defects in T cell development can be caused by abnormalities in the thymic epithelium, which provides a crucial supportive microenvironment for development of T cells^25^, or can be cell intrinsic to hematopoietic cells as, for instance, caused by mutations in genes required for T cell development. Because of crosstalk between developing thymocytes and the stromal cells, there often are secondary effects. To distinguish between direct and indirect effects, we made use of a widely accepted *in vitro* model system in which hematopoietic progenitor cells are cultured on OP9 stromal feeder layers that have been modified to express the Notch ligand DL1^26^, required to induce T cell development. Using this system, we took WT and *Morc3^MD41/MD41^* E14.5 fetal liver hematopoietic cells, co-cultured them for seven days on OP9-DL1 and performed flowcytometry analysis on DN cells (Fig. 4A). Comparing WT to *Morc3^MD41/MD41^* cultures, we observed a severe arrest in development at the DN1 stage (Fig. 4B-C), recapitulating the arrest previously observed *ex vivo* (Fig. 1). We further detected significantly reduced *Morc3* and *Tcf7* mRNA levels in *Morc3^MD41/MD41^* DN1 cells by RT-qPCR (Fig. 4D), consistent with our scRNA-seq result (Supplementary File 1). Flow cytometry also revealed increased myeloid and NK cell populations, but no B cells, in *Morc3^MD41/MD41^* cultures when compared to WT (Fig. 4E-F), in agreement with our ex vivo observations (Fig. 1C). Combined, these results indicate that the severe T cell developmental defects and expansion of non-T cell lineages observed are *Morc3^MD41/MD41^*hematopoietic cell intrinsic. Importantly, this defect is not caused by a deficiency in the fetal liver stem cell/progenitor cell compartment, as the fetal livers in *Morc3^MD41/MD41^* mice are similar in size, cellularity and contain stem cells at slightly increased numbers (Supplementary FigS1 and 7A-C).

**Figure 4.**
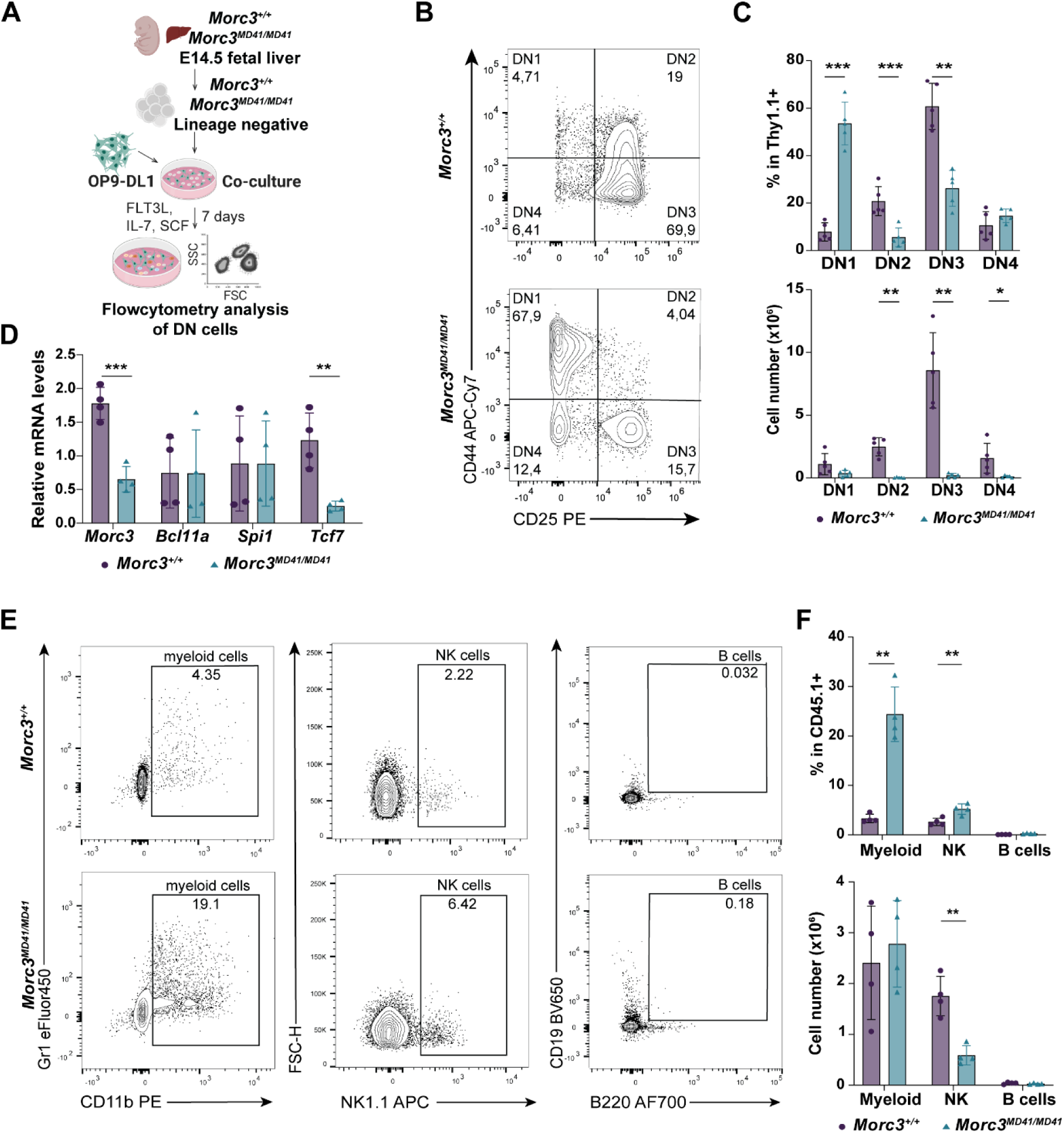
The T cell development phenotype in *Morc3^MD41/MD41^* homozygotes is cell intrinsic. (A) Schematic illustration of the differentiation of T cells from E14.5 fetal liver-derived lineage negative (Lin- : CD3-, CD4-, B220-, Gr1-, NK1.1-, Cd11c-, TER119-) cells co-cultured on OP9-DL1 cells. (B) Representative flowcytometry plots of *Morc3^+/+^* and *Morc3^MD41/MD41^* DN1, DN2, DN3 and DN4 cells generated from E14.5 fetal liver-derived lineage negative cells co-cultured on OP9-DL1 cells for 7 days. (C) (top) Percentage of DN1, DN2, DN3 and DN4 cells within Thy1.1+ cells (pre-gated on live, Lin-, CD45.1+) and (bottom) total cell number (n = 5 biological replicates per genotype, 3 independent experiments); data are presented as mean with dots/triangles as individual values; error bar represents SD. Statistical significance was determined using two-sided t test. (D) RT-qPCR analysis of relative *Morc3*, *Bcl11a*, *Spi1* and *Tcf7* mRNA levels in *Morc3^+/+^*and *Morc3^MD41/MD41^* sorted DN1 cells after 7 days of co-culture (normalized to β-actin; n = 4 biological replicates; data are presented as mean with dots/triangles as individual values; error bar represents SD. Statistical significance was determined using two-sided *t* test. (E) Representative flowcytometry plots of *Morc3^+/+^* and *Morc3^MD41/MD41^* myeloid (Gr1+, Cd11b+), NK (NK1.1+, Gr1-,Cd11b-) and B cells (CD19+, B220+, Gr1-,Cd11b-, NK1.1-, Thy1.1-) generated from E14.5 fetal liver-derived lineage negative cells co-cultured on OP9-DL1 cells for 7 days. All cells were pre-gated on single, live, CD45.1+. (F) (top) Percentage of myeloid, NK and B cells within CD45.1+ cells (pre-gated on single, live) and (bottom) total cell number (n = 4 biological replicates/genotype, 2 independent experiments); data are presented as mean with dots/triangle as individual values; error bar represents SD. Statistical significance was determined using two-sided *t* test. For all figures: *P≤0.05, **P≤0.01, ***P≤0.001, ****P ≤ 0.0001

### Chromatin accessibility analysis on *Morc3^+/+^* and *Morc3^MD41/MD41^*thymocytes shows decreased opening at T cell specific sites and enhanced accessibility at non-T lineage sites

The previously reported requirement for MORC3 in chromatin regulation, prompted us to investigate genome-wide chromatin accessibility in OP9-DL1 co-cultured, sorted DN1 cells using Assay for Transposase-Accessible Chromatin with high-throughput sequencing (ATAC-seq) (Fig. 5A). Comparing WT and *Morc3^MD41^* homozygous mutant cells, we found that loss of MORC3 had a mild genome-wide impact on chromatin accessibility. We found that 3.5% (n=881) of all peaks (n=25000) were significantly differentially accessible, among which 339 peaks were “more accessible” and 542 were “less accessible” in *Morc3^MD41^*homozygous DN1 cells (Fig. 5B-C and Supplementary FigS8A). Genomic annotation showed enrichment of *Morc3^MD41/MD41^* associated differentially accessible peaks in intergenic regions, introns and exons (Fig. 5D). In addition, promoters and UTRs (5’ and 3’) were affected, albeit to a lesser extent (Fig 5D). We did not find differences in chromatin accessibility at transposable elements (Supplementary FigS8B), a phenotype we previously saw in *Morc3^MD41^* homozygous mutant mESCs. Using HOMER, we next investigated whether differentially accessible peaks contain transcription factor (TF) binding motifs. We identified motifs for T cell commitment factors such as RUNX1 and BCL11B as significantly enriched sites with reduced accessibility, and motifs predicted to be targeted by EOMES and TFs belonging to the TBX family in more accessible regions in *Morc3^MD41/MD41^*DN1 cells (Fig. 5E). This is in agreement with the observed cell type restricted phenotype and indicates chromatin accessibility changes at regulatory sites specific for early T cell and NK cell developmental processes.

**Figure 5.**
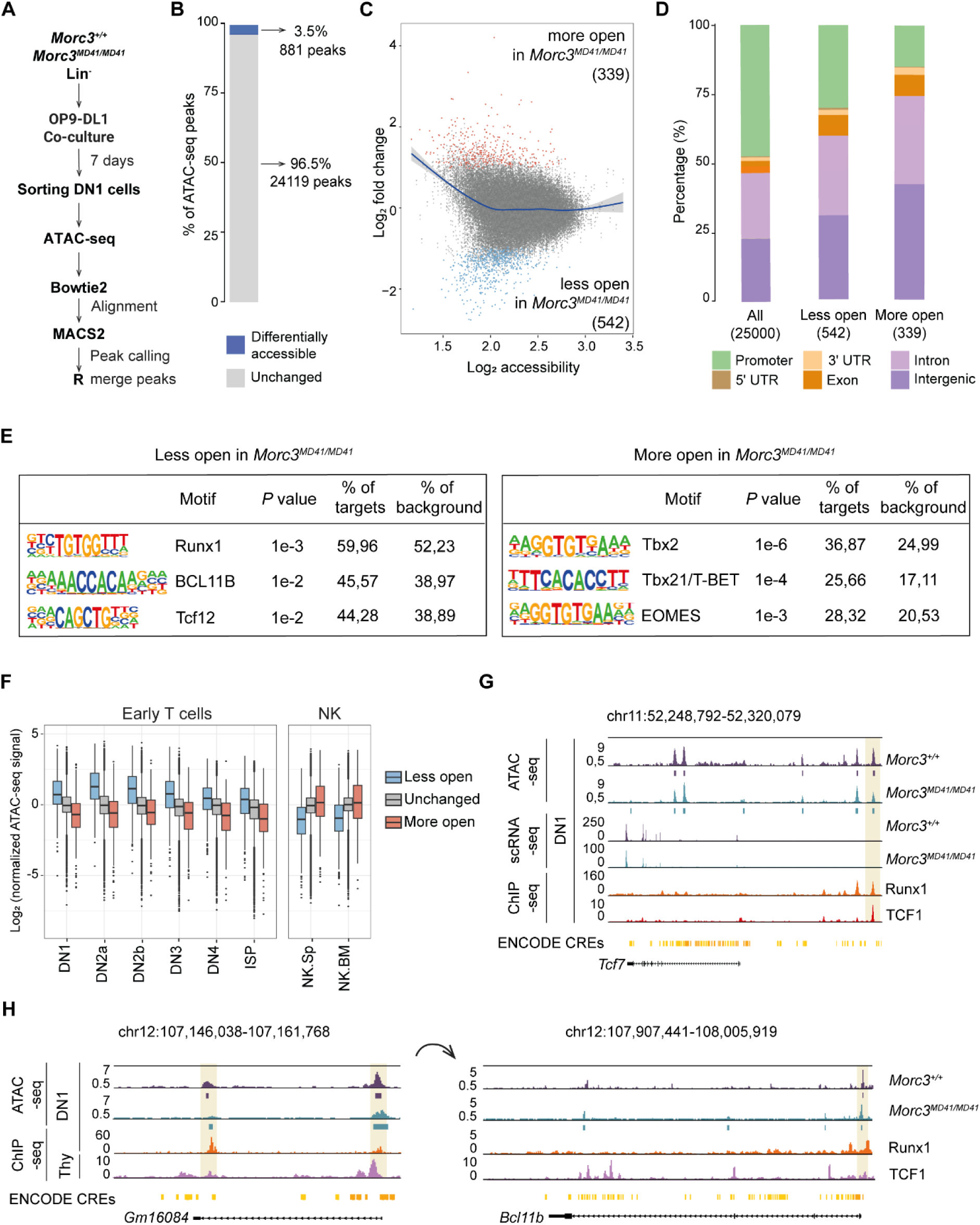
Chromatin accessibility analysis on *Morc3^+/+^* and *Morc3^MD41/MD41^* DN1 cells. (A) Schematic representation of the ATAC-seq experiment and analysis pipeline. (B) Stacked bar chart showing proportions of differentially accessible (881) and unchanged (24119) ATAC-seq peaks (calculated by Limma). (C) MA plot of changes in chromatin accessibility. Red indicates peaks with significant increased accessibility in *Morc3^MD41/MD41^* (p-adj < 0,05); blue indicates peaks with decreased accessibility in *Morc3^MD41/MD41^* (p-adj < 0,05). N = 3 biological replicates per genotype. (D) Stacked bar chart showing proportion of all, less and more open ATAC-seq peaks with respect to their genomic annotation. (E) Table showing motifs enriched at differentially accessible peaks, using HOMER analysis of known DNA sequence motifs. (F) Box plot depicting quantile normalized ATAC-seq signal of unchanged and differentially accessible peaks in ImmGen ATAC-seq dataset in early T cells and NK. Summits reported by ImmGen were matched to significant differentially accessible peaks by taking the summit closest to the peak centers of significant peaks. Boxes show the regions between the 25^th^ and 75^th^ percentiles. Upper whisker extends to highest value unless this extends beyond 1.5 times the interquartile range (IQR) (distance between first and third quartiles), in which case 1.5 times IQR is displayed. Lower whisker extends to lowest value, or 1.5 times IQR. Values beyond 1.5 times IQR are displayed as dots. The depicted cell types are: DN1; DN2a; DN2b; DN3; DN4, ISP, Immature Single Positive (from thymus); NK.Sp, Natural Killer (CD27+CD11b+ from spleen); NK.BM, Natural Killer (CD27+CD11b+ from bone marrow). (G) UCSC genome browser screenshot of the *Tcf7* locus. Representative tracks for ATAC-seq and scRNA-seq, published ChIP-seq (TCF1: GSM7549384; RUNX1: GSM2786857) from DN1 cells and ENCODE cis-regulatory elements (CREs) in orange boxes are shown. Yellow shading indicates representative differential ATAC-seq peaks overlapping with a known *Tcf7* enhancer element. (H) UCSC genome browser screenshot depicting (right) the *Bcl11b* locus and (left) a known enhancer. Representative tracks for ATAC-seq from DN1 cells, published ChIP-seq (TCF1 Thy (Thymocytes): GSM1133644; RUNX1 in DN1 cells: GSM2786857) and ENCODE cis-regulatory elements (CREs) in orange boxes are shown. Yellow shading indicates representative differential ATAC-seq peaks overlapping with (right) promoter and (left) enhancer element.

To test this, we matched *Morc3^MD41/MD41^* DN1 differentially accessible peaks with IMMGEN ATAC-seq datasets for various immune cell types and calculated a measure of specificity by normalizing the ATAC-seq signal of each peak by its average over the selected cell types. This analysis revealed that *Morc3^MD41/MD41^* DN1 less accessible peaks show higher specificity for DN thymocytes (Early T cells) than non-differentially accessible peaks (Fig. 5F and Supplementary FigS8C). Notably, these peaks also showed an even higher mean-normalized accessibility for DN2a cells (average of 3.45-fold more accessible in DN2a compared to 2.50-fold in DN1) (Fig. 5F), suggesting that loss of MORC3 prevents these genomic sites from opening. Furthermore, more accessible peaks in *Morc3^MD41/MD41^* DN cells matched loci associated with relatively higher chromatin accessibility in NK cells (Fig. 5F). This was supported by Gene Set Enrichment Analysis on a NK cell-specific gene set identified by the ImmGen consortium (coarse module 19) (Supplementary FigS8D).

Since *Morc3^MD41/MD41^* differentially accessible chromatin was enriched for intergenic regions (Fig. 5D), which can contain regulatory elements, we next examined whether these sites associated with putative ENCODE enhancers. This analysis showed that *Morc3^MD41/MD41^* DN1 less accessible peaks are significantly enriched for distal enhancer elements (224 out of 542 peaks; chi-square test P=0.003998) (Supplementary FigS8E), specific for DN1-3 thymocyte differentiation (Supplementary FigS8F). Representative genome browser screenshots of differentially accessible enhancers are shown for *Tcf7* and *Bcl11b* (Fig. 5G-H). Notably, *Tcf7* and *Bcl11b* are amongst several key T cell TF encoding genes identified as differentially expressed in our scRNA-seq (Supplementary File 1). Around one third of *Morc3^MD41/MD41^* DN1 more accessible peaks (133 out of 339 peaks) associated with ENCODE regulatory elements (Supplementary FigS8E). For instance, the *Crtam* gene, which plays a role in NK cell function, harbors two regulatory sites with increased chromatin accessibility in *Morc3^MD41^* homozygous DN1 cells (Supplementary FigS8G), and we observed an expansion of NK cells in *Morc3^MD41/MD41^* thymus (Fig. 3B-C) and in the *in vitro* culture (Fig. 4F). Collectively, these results indicate that T cell and NK cell regulatory regions are among the main targets of chromatin accessibility changes in *Morc3^MD41/MD41^* DN1 cells and that the dysregulated DN1 transcriptome was reflected at the chromatin level. These data further suggest that the *Morc3^MD41^*-associated T cell defect could be linked to the dysregulation of regulatory elements for multiple TFs critical for setting-up the T cell lineage program.

### Functional MORC3 ATPase and CW zinc finger domains are required for T cell development

To better understand how MORC3 regulates T cell lineage commitment, we next employed the OP9-DL1 co-culture system to investigate the role of the various MORC3 functional domains. We first determined, whether re-expressing WT MORC3 protein in *Morc3^MD41/MD41^* fetal liver stem/progenitor cells could restore normal T cell development and overcome the arrest at the DN1 stage. We used recombinant retroviruses to transduce WT and *Morc3^MD41^*homozygous, lineage negative, fetal liver cells with GFP-tagged full length MORC3 (Fig. 6A). While thymocyte development was arrested at the DN1 stage in the empty vector control, re-expression of WT MORC3 rescued thymocyte development in *Morc3^MD41^* homozygotes (Fig. 6B). This also underscored the cell intrinsic nature of the MORC3 deficiency.

**Figure 6.**
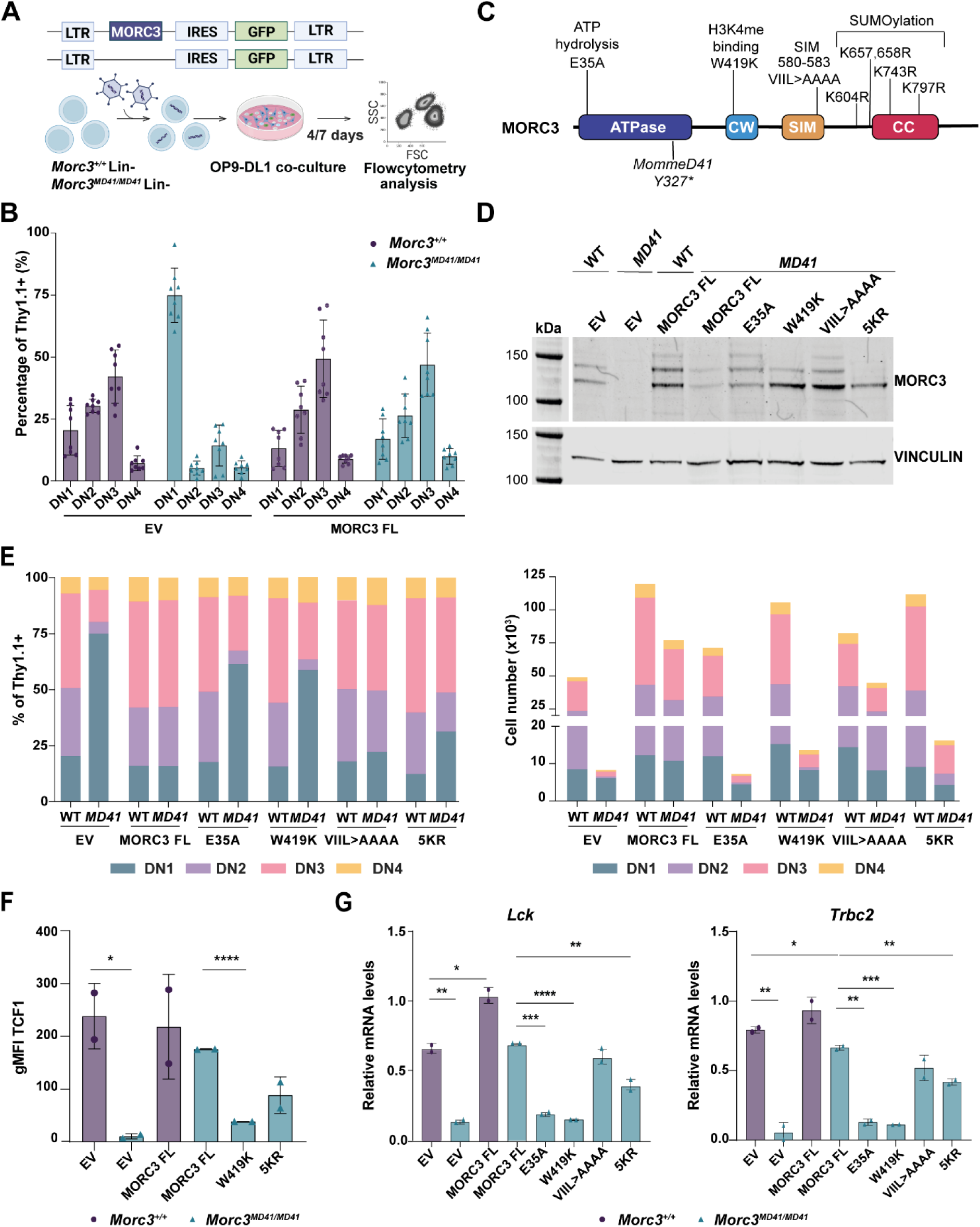
ATP hydrolysis and H3K4me binder functions of MORC3 contribute to T cell development. (A) Schematics of the rescue experiment with ectopic expression of full length MORC3 and empty vector (EV) in fetal liver-derived lineage negative (Lin-) cells and co-cultured on OP9-DL1. (B) Bar plot showing the percentage of DN1, DN2, DN3 and DN4 cells of *Morc3^+/+^* and *Morc3^MD41/MD41^*cells transduced with MORC3 full length (FL) virus and EV, on day 4 after OP9-DL1 co-culture. All cells are pre-gated on SSC-A/FSC-A, singles, live cells (7AAD-), Lin-, CD45.1+, GFP+ (only for EV transduced cells), Thy1.1+. (n = 8 per genotype, 4 independent experiments); data are presented as mean with dot/triangle as individual values; error bar represents SD. (C) Schematics of MORC3 protein domains and MORC3 mutants (E35A, W419K, VIIL>AAAA, 5KR). (D) Western Blot showing MORC3 expression in *Morc3^+/+^* and *Morc3^MD41/MD41^* lineage negative cells transduced cells with EV, MORC3 FL and mutants and sorted for GFP+ cells after 48 hours of culture. VINCULIN was used as loading control. (E) Stacked bar chart indicating the percentage (left) and total cell count (right) of DN1, DN2, DN3 and DN4 cells of *Morc3^+/+^* and *Morc3^MD41/MD41^* cells transduced with EV, MORC3 FL, MORC3 mutants (E35A, W419K, VIIL>AAAA, 5KR), on day 4 after OP9-DL1 co-culture. All cells are pre-gated on SSC-A/FSC-A, singles, live cells (7AAD-), Lin-, CD45.1+, GFP+ (only for GFP control transduced cells), Thy1.1+). Bars show mean from n = 8 EV, MORC3 FL; n = 6 W419K, 5KR; n = 4 VIIL>AAA, E35A, in 4 independent experiments. (F) Bar plot showing geometric mean fluorescent intensity (gMFI) of TCF1 expression in DN1 cells after 5 days of co-culture of *Morc3^+/+^* and *Morc3^MD41/MD41^* cells transduced with EV, MORC3 FL, W419K or 5KR MORC3 constructs. gMFI was calculated by subtracting gMFI of TCF1 expression in negative control (n = 2 biological replicates per genotype), data are presented as mean with dot/triangle as individual values; error bar represents SD. (G) RT-qPCR analysis of relative *Lck* and *Trbc2* mRNA levels in *Morc3^+/+^* and *Morc3^MD41/MD41^*cells after 4 days of OP9-DL1 co-culture. Data are normalized to *Ptprc*; n = 2 biological replicates per genotype; data are presented as mean with dot/triangle as individual values; error bar represents SD. For F and G panels, statistical significance was determined using two-sided *t* test. * P ≤ 0.05, ** P ≤ 0.01, *** P ≤ 0.001, **** P ≤ 0.0001.

Mouse MORC3 has a N-terminal ATPase required for dimerization and catalytic activity, a CW Zinc finger domain that recognizes methylated lysine 4 of H3 (H3K4me) and confers the chromatin reader properties, a SIM that allows interaction with other SUMOylated proteins and a C-terminal coiled-coil domain that harbors five SUMOylation sites^17,20,21,27,28^. In mESCs, MORC3 interacts with DAXX through SUMOylation and this MORC3-DAXX interaction is required for silencing of transposable elements^17^. To assess the importance of the different domains for thymocyte development, we generated a series of mutant constructs (Fig. 6C), for retroviral transduction of WT and *Morc3^MD41^* homozygous, lineage negative, fetal liver cells. Western blot showed that all variants were expressed at nearly endogenous levels, where the absence of SUMOylated MORC3 bands in the 5KR mutant functionally confirmed the mutations in the SUMOylation sites (Fig. 6D). We observed an almost complete rescue with WT MORC3 and a partial rescue when MORC3-SIM (VIIL>AAAA) and MORC3-SUMOylation (5KR) mutants were reintroduced, whereas the MORC3-CW-zinc finger domain (W419K) and MORC3-ATPase (E35A) mutants could not rescue the early thymocyte developmental block in *Morc3^MD41^* homozygotes when measured by flowcytometry after four or seven days of co-culture (Fig. 6E and Supplementary FigS9A-D). Importantly, re-expressing MORC3 not only phenotypically rescued thymocyte development, but also restored expression of key T cell markers such as TCF1 protein (Fig. 6F and Supplementary FigS9E-F) and mRNA levels of *Lck* and *Trbc2* (Fig. 6G and Supplementary FigS9G), suggesting that true DN2 and DN3 subpopulations were generated. Hence, in contrast to mESCs, the CW-zinc finger domain which mediates MORC3 binding to the active histone mark H3K4me is required for normal T cell development, whereas the contribution of MORC3 SUMOylation is mild.

### Expressing TCF1 or BCL11B in *Morc3^MD41/MD41^* cells rescues T cell development

We have previously shown that T cell development is driven by a transcription factor hierarchy starting with Notch signaling, which induces TCF1 expression that subsequently induces the expression of two target genes important for thymocyte development: the BCL11B TF that largely is responsible for further upregulation T cell specific genes and the GATA3 TF that mostly mediates the down regulation of non-T cell genes^7^. As chromatin accessibility of both *Tcf1* and *Bcl11b* regulatory elements and also mRNA levels of *Tcf7* and *Bcl11b* were decreased in *Morc3^MD41/MD41^* cells (Supplementary File 1), we performed retroviral overexpression studies of these TFs in a *Morc3^MD41/MD41^*background. This allows for an epistasis analysis to determine if MORC3 acts upstream of these TFs and in a dominant fashion in the regulation of T cell lineage commitment. Employing the OP9-DL1 co-culture system, we found that forced expression of either TCF1 or BCL11B could rescue T cell development in *Morc3^MD41/MD41^*cells.

TCF1 reconstitution resulted in a normal distribution over the DN subsets (Fig. 7A-B and Supplementary FigS10A-B), and BCL11B even stimulated development of later stages such as DN4 thymocytes (Fig. 7C-D), probably because it normally is expressed slightly later than TCF1. These data formally show that MORC3 acts upstream of TCF1 and BCL11B and that dysregulation of these lineage specific transcription factors is a main reason for the T cell developmental arrests observed due to MORC3 loss of function.

**Figure 7.**
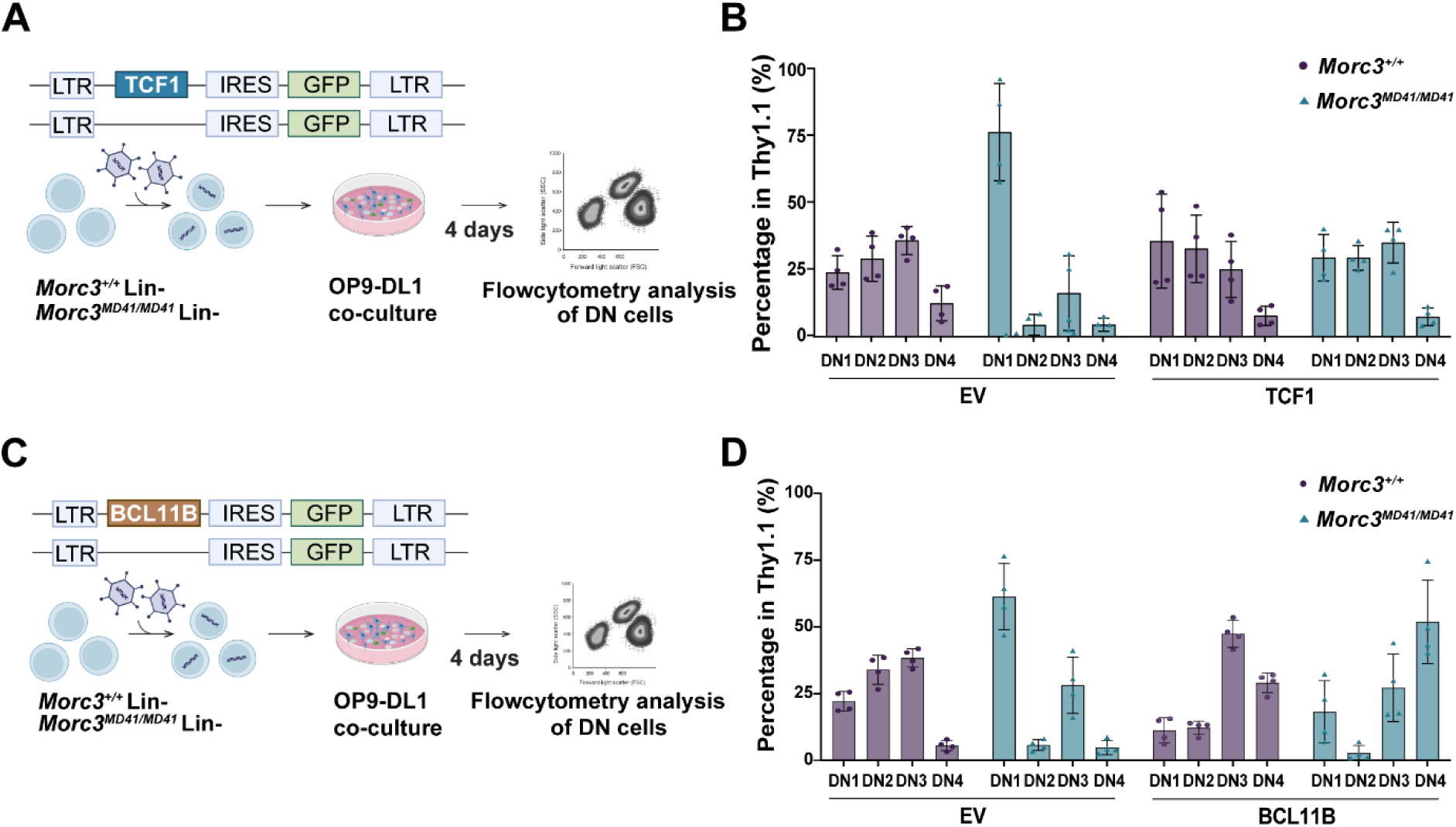
Ectopic expression of TCF1 and BCL11B rescues T cell development in *Morc3^MD41/MD41^* cells. (A) Schematics of the rescue experiment with ectopic expression of TCF1 and empty vector (EV) in fetal liver-derived lineage negative (Lin-) cells and cultured on OP9-DL1. (B) Bar plot showing the percentage of DN1, DN2, DN3 and DN4 cells of *Morc3^+/+^* and *Morc3^MD41/MD41^*cells transduced with EV or TCF1, on day 4 after OP9-DL1 co-culture. All cells are pre-gated on SSC-A/FSC-A, singles, Lin-, CD45.1+, Thy1.1+. n = 4 biological replicates per genotype; 2 independent experiments. Data are presented as mean with dot/triangle as individual values; error bar represents SD. (C) Schematics of the rescue experiment with ectopic expression of BCL11B and EV in fetal liver-derived lineage negative (Lin-) cells and cultured on OP9-DL1. (D) Bar plot showing the percentage of DN1, DN2, DN3 and DN4 cells of *Morc3^+/+^* and *Morc3^MD41/MD41^* cells transduced with EV or BCL11B, on day 4 after OP9-DL1 co-culture. All cells are pre-gated on SSC-A/FSC-A, singles, Lin-, CD45.1+, GFP+, Thy1.1+. n = 4 biological replicates/genotype; 2 independent experiments. Data are presented as mean with dot/triangle as individual values; error bar represents SD.

## Discussion

Developmental lineage decisions are regulated by the formation of a cell type–specific epigenetic landscape and a network of transcriptional activators and repressors. As binding of most TFs is influenced by chromatin state, the cell type-specific epigenetic landscape needs to be established before or concurrently with the expression of lineage-specifying TFs. However, while the functions of lineage-determining TF and their targets are well understood, the epigenetic factors that participate in these processes have remained largely elusive.

Here, we report that loss of the chromatin reader MORC3 has a severe effect on thymic T cell development at the most immature stages in vivo. Most thymocytes are arrested at the DN1 stage and while some cells can proceed a bit further, development is fully blocked at DN3, coinciding with T cell lineage commitment. Indeed, development of NK cells and various myeloid lineages that can also develop in the thymus proceeds normally, while mature T cells are completely absent in *Morc3^MD41/MD41^*homozygotes. This phenotype is reminiscent of mice with targeted mutations in *Tcf7*, *Notch1*^24^ and *E2A*^29^. However, Notch1 mutations in lymphocytes also lead to an accumulation of B cells, and Notch target genes such as *Hes1* are unaffected in *Morc3^MD41/MD41^* thymocytes. *E2A* and *Bcl11b* mutant mice show an arrest in T cell development a bit later than what we observe in MORC3 deficient cells, while TCF1 deficient stem cells when forced into the T cell lineage block completely at DN1^30,31^. Collectively, a large part of the thymic phenotype in *Morc3^MD41^* homozygotes can be explained by compromised TCF1 levels. Indeed, forced expression of TCF1 rescues T cell development from *Morc3^MD41/MD41^* thymocytes.

The block in T cell development and skewing towards the myeloid lineage could be rescued by reintroducing WT MORC3 protein. Protein domain analysis indicated that MORC3 facilitates these effects through its CW Zinc-finger, which binds methylated forms of the active histone mark H3K4^20,21^, and its ATPase domain. This is in marked contrast to transposable element silencing by MORC3 in mESCs, which involves MORC3 localization to the repressive histone mark H3K9me3^16,17^, MORC3 SUMOylation and DAXX interaction^17^. Our data reveal, for the first time, a functional requirement for MORC3’s CW zinc-finger domain and suggest cell type-specific roles for MORC3, which could be conferred by MORC3 being part of different protein complexes.

ATAC-seq indicated that MORC3 loss led to altered chromatin accessibility in DN1 cells, where T cell specific regulatory sites are more closed and non-T cell regulatory sites showed more open chromatin. And while Notch signaling is largely unaffected, the expression of important TFs for T cell development such as TCF1 and BCL11b is reduced in *Morc3^MD41^* homozygotes. Overexpression of TCF1 and BCL11B rescued T cell development of *Morc3^MD41/MD41^* progenitors, indicating that MORC3 acts upstream of these TFs. Our data suggest that MORC3 influences gene expression via chromatin accessibility and it is possible that MORC3, by binding to H3K4me, helps to maintain the proper chromatin landscape for T cell TFs to bind and exert their function. However, we cannot currently link genomic sites with altered chromatin accessibility to MORC3 occupancy in DN1 cells directly. Alternative scenarios are therefore possible. For instance, since *Morc3^MD41/MD41^*-associated differentially accessible peaks are enriched for DNA motifs that are recognized by TFs, it is possible that MORC3 gets recruited to these sites through direct interaction with TFs, such as TCF1.

In sum, these findings advance our understanding of the cross-talk between epigenetic regulators and transcription factors specific for T cell development. While our studies are currently confined to mouse models, we note that MORC3 is similarly expressed in human thymocytes^32^. It will be of interest to explore a potential role for MORC3 in human T cell development and human T-cell acute lymphoblastic leukemia, the malignant counterpart of thymocytes, in future studies.

## Material and Methods

### Ethics statement

All procedures involving animals were approved by the Animal Ethics Committee of the Leiden University Medical Centre and by the Commission Biotechnology in animals of the Dutch Ministry of Agriculture.

### Mice

The *MommeD41* line has been previously generated on the FVB/NJ Line3 background^16^ and was maintained as heterozygous. Mice were housed under standard conditions in the pathogen-free Animal Care Facility of the LUMC, with free access to food and water at all times. For tissue and embryo collection, mice were euthanized by cervical dislocation. Natural timed matings were set up and the morning of the vaginal plug was designated Embryonic day 0.5 (E0.5).

### Genotyping

Genomic DNA (gDNA) was extracted using the salt-extraction method. Tissues were lysed and digested in cell lysis buffer (50 mM Tris-HCl (pH 8), 4 mM EDTA (pH 8), 2% SDS) with Proteinase K (390973P, VWR) and incubated at 55°C overnight. RNAseA (EN0531, Thermo Fisher Scientific) treatment was performed at 37°C for 1 hour, followed by the addition of saturated sodium chloride. gDNA was precipitated with isopropanol and washed with 70% ethanol. The extracted gDNA was dissolved in water. DreamTaq Polymerase (EP0705, Thermo Fisher Scientific) was used for genotyping and primer sequences are provided in Supplementary File 2. Before Sanger sequencing, PCR products were cleaned up with Exonuclease I (70073X 5000 UN, Affymetrix) and FastAp Thermosensitive Alkaline Phosphatase (EF0654, Thermo Fisher Scientific) and incubated at 37°C for 1 hour and at 80°C for 15 minutes.

### Protein isolation

Tissues were lysed in Protein Lysis buffer (20mM Triethanolamine, 150mM NaCl, 1% Sodium Deoxycholate, 1% SDS, 1% Triton X-100) with Protease Inhibitor Cocktail (27368400, Roche) and Phosphatase Inhibitor Cocktail (4906837001, Sigma) on ice using glass beads (0.5mm, GB05, Bio-Connect) and bullet blender Storm24 (Next Advances; intensity 10, 30 seconds). A BCA kit (23225, Thermo Scientific) was used to measure protein concentration. Sorted cells were lysed in RIPA buffer (0,1% SDS, 1% NP-40, 150mM NaCl, 5mM EDTA, 0,5% Sodium Deoxycholate, 20mM Tris-HCl) with Protease Inhibitor Cocktail (27368400; Roche) on ice by using resuspension.

### Western blot

Total cell extracts were loaded on a NuPAGE Bis-Tris gel (4–12%, NP0321; Thermo) or Criterion XT Bis-Tris (4-12%, 3450124, Bio-Rad), and transferred to a Nitrocellulose Blotting Membrane (10600016; Life Sciences). The following primary antibodies were used: MORC3 (100-401-N96S; Rockland; 1:1000) and VINCULIN (V9131; Merck, 1:2000). Donkey anti-Rabbit 800CW (926-32213; Li-Cor, 1:5000) and Donkey anti-mouse 680RD (926-68072; Li-Cor, 1:5000) were used as secondary antibodies. Membranes were analyzed on Odyssey (Westburg).

### Single cell suspensions

Tissues were pressed through a 35 μm nylon cell strainer (352235, Corning) for thymus and 70 μm nylon cell strainer (352350, Corning) for fetal liver. Cells were resuspended in IMDM (Iscove’s Modified Dulbecco’s Medium, Capricorn) supplemented with 2,5% FBS (fetal bovine serum, Bodinco) and 1% penicillin/streptomycin (15070-063, Gibco). Thymocytes were used immediately for flow cytometry assays and fetal liver cells were stored in liquid nitrogen until further application.

### Flowcytometry analysis and cell sorting

Cells were stained with appropriate fluorochrome-conjugated antibodies and biotinylated antibodies (Supplementary File 2) and incubated for 30 minutes at 4°C in the dark in FACS buffer containing phosphate buffered saline (PBS, 3623140, Braun) with 2% bovine serum albumin (BSA, A9647, Sigma) and 0.1% sodium azide. A second labelling step was conducted using fluorochrome-conjugated streptavidin (Supplementary File 2) and 7-AAD (559925, BD Biosciences) was used as a live/dead marker. Gating strategy is provided in Supplementary FigS11.

### Apoptosis and proliferation assays, TCF1 intracellular staining

For detection of apoptotic cells, cells were stained with Annexin V (563974, BD Biosciences) and 7-AAD (559925, BD Biosciences) in diluted Annexin V Binding Buffer, 10X concentrate (556454, BD) after cell surface staining. For proliferation assay and TCF1 expression measurement, the intracellular staining was performed using eBioscience™ Foxp3 / Transcription Factor Staining Buffer Set (00-5523-00, Thermo Fisher Scientific) after cell surface staining. Cells were measured on Aurora (Cytek), FACSCanto II (BD Biosciences) and LSR Fortessa X-20 (BD Biosciences). Cell sorting was performed on FACSAria II (BD Biosciences). Data were analyzed using FlowJo software (Tree Star).

### Single cell RNA-sequencing (scRNA-seq)

Single cell suspensions were freshly prepared from embryonic thymus. Dissections were done blind to genotype and one homozygous and one control (heterozygous) thymus were selected based on organ size and total cell count. Freshly isolated thymocytes were stained with eFluor450-conjugated CD45 (48-0451-82, eBioscience) and 7-AAD (559925, BD Biosciences) in FACS buffer for 30 minutes at 4°C. From each sample, 20.000 live leukocytes (CD45+,7AAD-) were sorted by FACSAria II cell sorter (BD Biosciences) and used for library preparation. scRNA-seq libraries were prepared using the 10X Chromium Next GEM Single Cell 3’ v3 kit following the manufacturer’s instructions. One library per sample was prepared and sequenced on an Illumina NovaSeq 6000 aiming for 2500 reads per cell.

### scRNA-seq analysis

UMI (Unique Molecular Identifiers) containing reads were mapped and counted with Cell Ranger 3.1.0 using the mouse genome Ensembl gene model file Mus_Musculus.GRCm38.gtf. For the *Morc3^MD41/+^*, the mean reads per cells was 35,180. For the *Morc3^MD41/MD41^*, the mean reads per cell was 38,427. The outputs of Cell Ranger were then processed individually using Seurat v3.1.4^33^ to filter out low-quality cells (Feature > 400, Features < 6000, Percentage of Mitochondrial genes < 5 %). Filtered libraries were normalized using the method « LogNormalize » and scale factor = 10000. Cell cycle regression was performed using the ensemble database (mmusculus_gene_ensembl dataset) with package bioMart 2.58.0. cc.genes.updated.2019$s.genes and cc.genes.updated.2019$g2m.genes. After normalization, data was clustered using the top 14 dimensions. To illustrate the differences between the clusters, principal component analysis was plotted with UMAP using “RunUMAP” function. Cluster identities were assigned based on the presence or absence of previously reported marker genes^1,34^. Subsets of the datasets, including only DN1, DN2a and DN3 cell population, were created for the further investigation of the differences between the groups. These subsets were merged with “merge” command, normalized and integrated into one dataset with “IntegrateData” command. Differential gene expression analysis was performed using the Seurat function “FindMarkers” by comparing similar cell annotation between *Morc3^MD41/+^* and *Morc3^MD41/MD41^* after integration of the datasets. To avoid the influence of the sex chromosome genes, these genes were excluded. The gene list was extracted from ensemble database “mmusculus_gene_ensemb” using a biomaRt package 2.58.0. Volcano plots were made using VolcaNoseR^35^. Gene ontology analysis was performed using PANTHER^36^.

### RNA isolation and RT-qPCR

Total RNA was extracted using QIAzol (5346994, QIAGEN) or RNeasy Micro Kit (74004, QIAGEN) for small numbers of sorted cells (25,000-50,000 cells), according to manufacturer’s recommendations. 150-500ng of RNA was reverse transcribed with RevertAid H Minus First Strand cDNA Synthesis Kit (K1632, Thermo Fisher Scientific). RT-qPCR was carried out with iQ™ SYBR® Green Supermix (1708887, Bio-Rad) on C1000TM Thermal cycler (Bio-Rad). Primers sequences are listed in Supplementary File 2.

### RNA-sequencing (RNA-seq) analysis

Total RNA was isolated as described above and RNA-seq was performed at BGI. Libraries were sequenced with 100 bp pair-end (PE) reads on a BGISEQ (DNBseq) platform. Quality assessment of the raw sequencing reads was done using FastQC v0.11.2 (http://www.bioinformatics.babraham.ac.uk/projects/fastqc). Adapters were removed by TrimGalore v0.4.5 (https://www.bioinformatics.babraham.ac.uk/projects/trim_galore/) using default parameters for paired-end Illumina reads, after which, quality filtering was performed by the same software. Reads smaller than 20bps and those with an error rate (TrimGalore option "-e") higher than 0.1 were discarded, after which a final quality assessment of the filtered reads was done with FastQC to identify possible biases left after filtering. The remaining reads were mapped to the mouse reference genome (build mm10) using STAR aligner v2.5.1^37^ and default parameters with the following exceptions: “– outputMultimapperOrder random” and “–twopassMode basic”. Before mRNA quantification, duplicated reads were marked with Picard tools v2.17 (http://broadinstitute.github.io/picard/). Quantification was done by HTSeq-count v0.91^38^, using the GENCODE MV16 annotation with the option “–stranded no”. Statistical analysis was done using DESeq2 v1.2.0^39^ (R package). Heatmap was created using ClustVis^40^; Volcano plot was made using VolcaNoseR^35^.

### In vitro OP9-DL1 co-culture

For in vitro differentiation of T cells, E14.5 fetal liver-derived hematopoietic progenitors were used as input. Single-cell suspensions were stained for lineage markers using biotin-conjugated antibodies (TER-119, B220, CD3, CD4, NK1.1, CD11c, Gr1), incubated with streptavidin-coated magnetic beads (130-048-101, Miltenyi Biotec) and passed through a magnetic column (130-042-401, Miltenyi Biotec) to deplete lineage-positive cells and enrich for hematopoietic progenitors. The lineage-negative cell enrichment was assessed by flow cytometry using antibodies against lineage markers, Sca1, and c-kit (data not shown). Lineage-negative cells were cultured overnight in StemSpan™ SFEM (09600, STEMCELL Technologies) supplemented with 10 ng/ml of recombinant mouse thrombopoietin (488-TO/CF, R&D Systems), 50 ng/ml of recombinant mouse Flt3-ligand (rmFlt3L) (427-FL, R&D Systems), and 100 ng/ml of recombinant mouse stem cell factor (rmSCF) (455-MC, R&D Systems). The following day, the cells were cultured on OP9-DL1 monolayer^26^ using αMEM (Minimum Essential Medium Eagle - alpha modification) (MEMA-XRXA, Capricorn), 10% FBS (Serana), 1% penicillin/streptomycin (15070-063, Gibco), and GlutaMAX (35050061, Gibco) medium complemented with 50 ng/ml of rmFlt3L, 10 ng/ml of rmSCF, 10 ng/ml of recombinant mouse interleukin-7 (rmIL-7) (407-ML, R&D Systems), and 50 μM β-mercaptoethanol (β-ME) (M3148-25ML, Sigma-Aldrich). After 2 and 5 days of culture, complete medium was refreshed. On day 4/7, cultured cells were harvested and analyzed by flow cytometry or sorted.

### Assay for transposase-accessible chromatin with sequencing (ATAC-seq)

ATAC-seq libraries were prepared as previously described^41^. DN1 cells (7-AAD-, CD45.1+, Lin-, Thy1.1+, CD44+, CD25-) derived from OP9-DL1 co-culture system were sorted in FBS and washed in cold PBS. 50.000 cells were resuspended in 50 µL transposition mix (Nextera, Illumina) and incubated for 30 min at 37 °C. Libraries were amplified by PCR with barcoded Nextera primers. Samples were sequenced at Macrogen on a NovaSeq6000 with 150bp pair-end (PE) reads.

### ATAC-seq analysis

After preprocessing with Trimmomatic (standard settings - seed mismatch 2, palindrome clip threshold 30, clip threshold 10, lead and train minimum trim quality 3, window size 4, average minimum quality of 15, minimum read length 20)^42^, reads were aligned to the mm10 version of the mouse genome using bowtie2 with options *“--very-sensitive --maxins 2000 --minins 0*”^43^. Sambamba was used to select only properly paired reads mapped to positions on chromosomes 1 to 19 (excluding sex chromosomes) and deduplicate alignments^44^. *MACS2* was used to call peaks for each sample using *“--nomodel --bdg -q 0.05*” settings^45^. The called overlapping peaks were merged in *R* using the reduce and *findOverlaps* from *GenomicRanges* and *GenomicAlignments*. Only peaks that were called in all three replicates of either WT or *Morc3^MD41/MD41^* samples were included in further analysis. BigWig files with RPKM were created using *bedtools genomecov* function^46^. Coverage plots and heat maps were generated with *deeptools* (bin size of 10bp) and smoothing the signal using a running mean on 11 consecutive bins^47^. Repeatmasker (Repeat Library 20140131) was used to define genomic regions with repeats. Top 1% highest scoring elements were used for LINE, LTR and SINE elements (31958, 173241 and 16244 repeats respectively). The statistically significant differentially accessible peaks were calculated using *limma* package (adjusted P value < 0.05) with gender of the mice as blocking factor^48^. The genomic annotations of ATAC-seq peaks were defined using *annotatePeak* function of ChIPseeker^49^. Motif analysis on the differentially accessible regions was performed using HOMER^50^, with the total set of peaks as background. Gene-set enrichment analysis (GSEA) was performed with *Fgsea* on ImmGen Coarse Module 19 (NK cell associated genes) (https://www.immgen.org/ModsRegs/modules.html) by pre-ranking genes according to signed p-values. To pair all peaks from this analysis with ATAC-seq peaks of immune cell populations generated by the Immunological Genome Project (ImmGen) database^51^, DiffBind^52^, was first used to calculate the summits of ATAC-seq peaks in WT and *Morc3^MD41/MD41^* DN1 cells. Nearest summits within 500bp were used to pair ATAC-seq peak summits from the ImmGen datasets. To calculate mean-normalized ATAC-seq values for each peak, the quantile normalized ATAC-seq signals published by^51^ were divided by their mean over all cell types. Annotations for candidate Cis Regulatory Elements (cCREs) were obtained from ENCODE (accession ENCSR412JPD). *FindOverlaps* from *GenomicRanges* was used to annotate peaks with cCRE classifications. In case of multiple annotations, the annotation with the largest overlap was used. cCRE classifications were simplified to the following: promoter like signatures (PLS), promoter-proximal enhancer like signatures (pELS), enhancer like signatures (ELS), regions with DNAse signal as well as H3K4me3 (DNAse-H3K4me3) and regions with DNAse signal only (DNase). Chi-squared test was performed on a two-by-two contingency table comparing less accessible peaks to unchanged peaks and comparing dELS annotated peaks to all other peaks. This was done with the *chisq.test* function in *R* using 2000 monte-carlo simulations (*simulate.p.value=True*).

### Cloning and generation of MORC3 mutants

In this study, full length *Morc3* cDNA was synthesized from RNA of mouse embryonic stem cells using polyA primers and the RevertAid First Strand cDNA Synthesis Kit. Gibson Assembly (GA, Gibson Assembly® Cloning Kit, NEB, E5510S) was used to clone *Morc3* cDNA into pCDNA_CMV_3Ty1 vector. The resulting pCDNA_CMV_3Ty1_Morc3 FL vector served as a template for in vitro mutagenesis to generate MORC3 mutants. For the five SUMOylation sites mutants (5KR), PCR fragments were generated from the *Morc3* FL clone using HiFi PCR (Takara PrimeSTAR GXL, R050A), with primers containing the desired KR mutant sequence and 40 nucleotides of 5’ overlap with adjacent fragments. Overlap PCR was performed, and the resulting 1kb fragment was cloned back in pCDNA_CMV_3Ty1_Morc3 FL vector. MORC3 mutants E35A, VIIL>AAAA, and W419K were generated by site-directed mutagenesis, using overlapping primers, with one of them containing the desired mutation. *Morc3* FL cDNA was cloned into the LZRS_IRES_eGFP (Addgene, 21961) vector by GA. The resulting LZRS_Morc3 FL_IRES_eGFP clone underwent partial HindIII and complete XhoI digestion, with a 2.6 kb fragment removed. *Morc3* cDNA containing with VIIL>AAAA, W419K, or 5KR mutations was removed from the pCDNA plasmid and cloned into the LZRS vector. For the E35A mutation, a 5’ 636 nt BamHI – EcoRI fragment was sub-cloned from LZRS_Morc3 FL_IRES_eGFP to pBS, introducing the mutation via in vitro mutagenesis. The resulting vector was cloned back into the BamHI-EcoRI-digested LZRS_Morc3 FL_IRES_eGFP vector. All constructs were confirmed by Sanger sequencing. All primers used during the cloning strategies are listed in Supplementary File 2.

### Retrovirus production

For retroviral packaging of GFP control, MORC3-FL and MORC3 mutant plasmids the Phoenix ecotropic cell line (CRL-3214, ATCC) was used. The cell line was checked regularly by measuring mouse CD8 expression (gag-pol) by flowcytometry and selected using Hygromycin B (10843555001, Roche) and Diphtheria Toxin (BML-G135-0001, Enzo). The day before transfection, 4×10^6^ cells were seeded in complete IMDM medium, containing 10% FCS, 1% penicillin/streptomycin, GlutaMAX. The retroviral constructs were transiently transfected using Xtremegene HP transfection reagent (6366236001, Sigma), following the manufacturer’s instructions. Media was changed 24 hours post-transfection and viral supernatants were harvested 24 hours later and stored at -80C until further use. TCF1 (p45) long isoform cloned in LZRS-eGFP backbone retrovirus was harvested from a stable Phoenix ecotropic cell line described previously^53^.

### Retroviral transduction and OP9-DL1 co-culture

E14.5 fetal liver-derived lineage negative cells were transduced using RetroNectin (T100B, Takara Bio Inc.) coated wells following the manufacturer’s instructions. Briefly, non-tissue culture plates (351172, CORNING) were coated with RetroNectin overnight at 4°C. RetroNectin was removed, and viral supernatant was added, followed by centrifugation at 1500xg for 2 hours at 32°C. Unbound retrovirus was discarded, and lineage negative cells were added to the virus-coated plates. The cells were cultured in StemSpan™ SFEM medium supplemented with rmTPO (10 ng/ml), rmFlt3L (50 ng/ml), and rmSCF (100 ng/ml). Transduction was performed overnight at 37°C. For testing transduction efficiency, GFP+ cells were FACS sorted 72 hours post-transduction and lysed for protein isolation (see above). For in vitro T cell differentiation, after overnight transduction, cells were seeded onto the OP9-DL1 cell line, using the above-described method. After 4 and 7 days of co-culture, cells were harvested to assess T cell development using flow cytometry analysis.

### Statistical analyses

Statistical significance was determined using unpaired Student’s t-test or two-way ANOVA as indicated in figure legends. All statistical analysis were performed in Prism 9 GraphPad software (version 9.3.1).

### Public datasets

ChIP-seq data for H3K4me1 in DN2 (GSM241836), EOMES (GSM5134535), Tbx21/T-BET (GSM5134533), H3K4me1 (GSM1441299) in NK were obtained from CistromeDB (Liu, 2011). BigWig files from CUT&RUN for TCF1 in DN1 were obtained from GSM7549384. ChIP-seq for Runx1 in DN1 (GSM2786857), TCF1 in thymocytes (GSM1133644) and H3K4me2 in DN1 (GSM774273) were downloaded and analyzed. Briefly, after preprocessing with TrimGalore to remove low-quality reads, reads were aligned to the mouse genome (mm10) using bowtie2 v2.3.4.2^43^ with parameters -very sensitive. To generate tracks, deepTools^47^ version 3.1.3 was used.

## Acknowledgements

We thank all members of the Staal and Daxinger laboratories for scientific discussions. We thank the animal caretakers for maintaining the mouse lines. We thank the Leiden Genome Technology Center for help with library prep and sequencing services and the LUMC Flowcytometry Core facility. Some illustrations were made using BioRender. Work in the Staal laboratory was supported in part by a ZonMW E-RARE grant (40-419000-98-020) and EU H2020 grant RECOMB (755170-2) and has received funding from the European Union Horizon 2020 research and innovation program as well as from reNEW, the Novo Nordisk Foundation for Stem Cell Research (NNF21CC0073729). In the Daxinger laboratory, this work was in part supported by a LUMC Fellowship, NWO ALW-Open (ALWOP.358) and NWO-Aspasia (015.014.035/6573) grants to LD.

## Author contributions

V.D.C. contributed to study design, performed experiments, analyzed and interpreted data. M.V. contributed to study design, performed experiments, analyzed and interpreted data. J.C. analyzed and interpreted scRNA-seq and ATAC-seq data. L.G.P. performed experiments, analyzed and interpreted data. G.N. and K.K.D.V. performed experiments. C.L. analyzed ATAC-seq data. A.K. analyzed scRNA-seq data. Z.Z. interpreted results. C.B. generated plasmids. S.G.V. performed experiments. K.C.B and K.P.O. interpreted results. S.E.J. interpreted results and contributed to manuscript writing. F.J.T.S. and L.D. designed and conceived the study, secured funding, supervised the study and wrote the manuscript. All authors contributed to manuscript writing and have read and agreed to the manuscript.

## Data availability

The RNA-seq, scRNA-seq and ATAC-seq data generated in this study have been deposited in the NCBI Gene Expression Omnibus (GEO).

