## Supplementary Figures for "The epigenetic reader MORC3 is required for T cell development in the thymus"

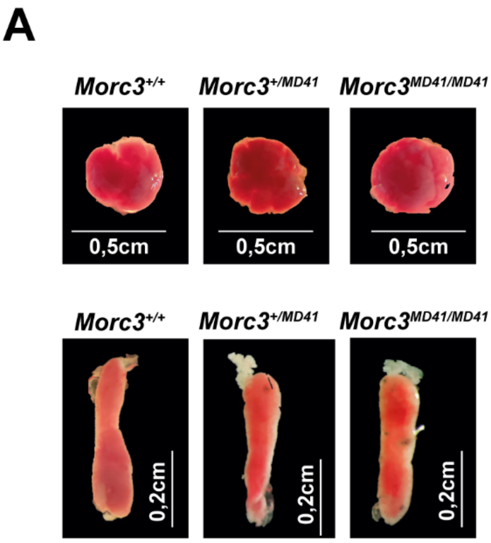


**Supplementary Figure S1.** ***Morc3$^{MD41/MD41}$*** **fetal liver and spleen have a normal morphology.**

(A) (top) Representative pictures of E14.5 fetal liver from *Morc3$^{+/+}$*, *Morc3$^{+/MD41}$* and *Morc3$^{MD41/MD41}$* embryos. (bottom) Representative pictures of E18.5 spleen from *Morc3$^{+/+}$*, *Morc3$^{+/MD41}$* and *Morc3$^{MD41/MD41}$* embryos.

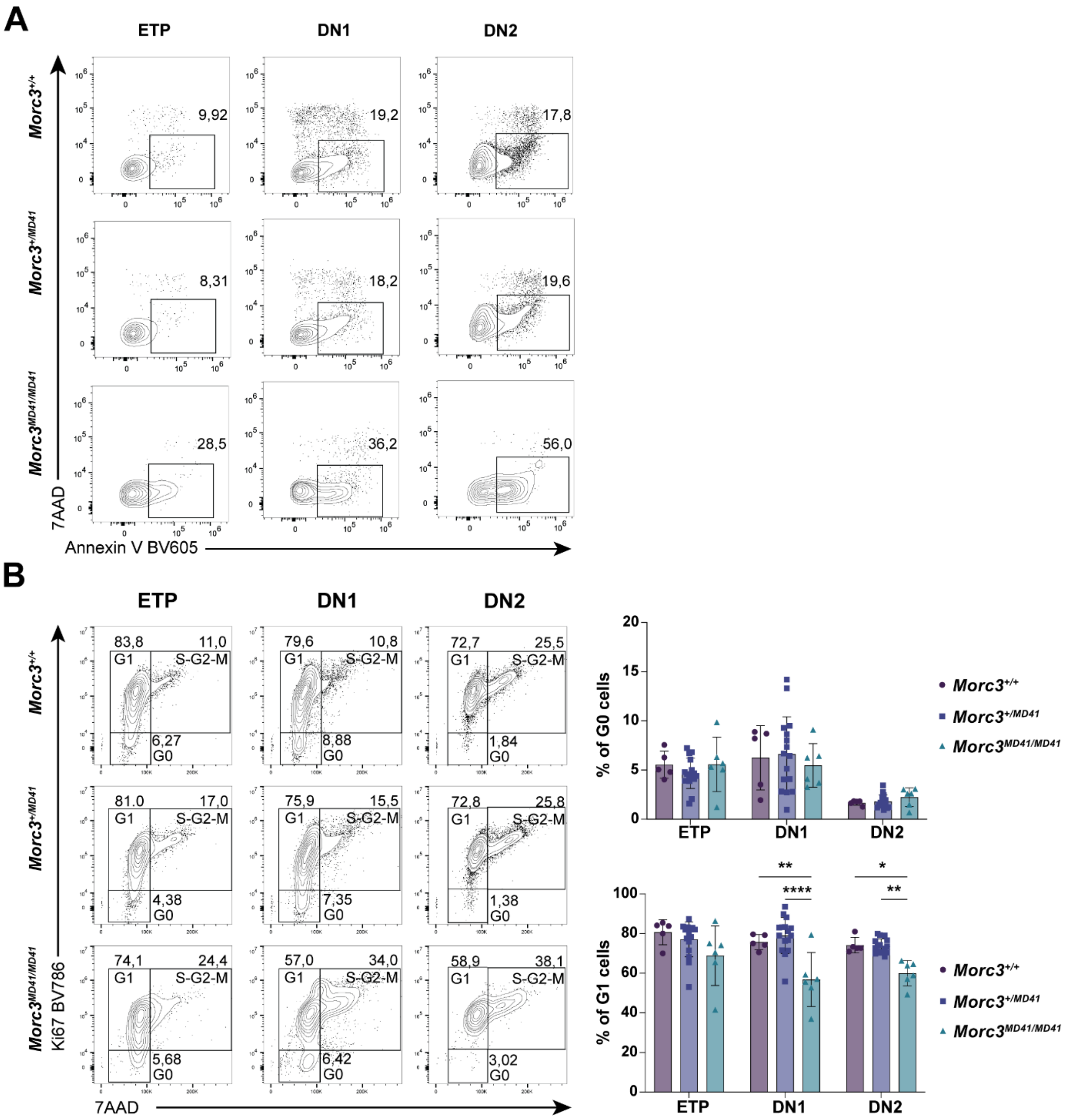


**Supplementary Figure S2. Apoptosis and cell cycle analysis in early T cells.**

(A) Representative flowcytometry plots of apoptosis analysis (apoptotic cells: Annexin V+, 7AAD-) in ETP, DN1 and DN2 cells in *Morc3*$^{+/+}$, *Morc3*$^{+/MD41}$ and *Morc3*$^{MD41/MD41}$ E18.5 thymus. (n = 8 *Morc3*$^{+/+}$, n = 8 *Morc3*$^{+/MD41}$, n = 5 *Morc3*$^{MD41/MD41}$, 3 independent experiments).

(B) (left) Representative flowcytometry plots of G0 (Ki67-, 7AAD-), G1 (Ki67+, 7AAD-), G2,S,M cells (Ki67+, 7AAD+) in ETP, DN1 and DN2 cells in *Morc3*$^{+/+}$, *Morc3*$^{+/MD41}$ and *Morc3*$^{MD41/MD41}$. (right) Percentages of G0 cells (top) and of G1 (bottom) within ETP, DN1 and DN2 cells in *Morc3*$^{+/+}$, *Morc3*$^{+/MD41}$ and *Morc3*$^{MD41/MD41}$ E18.5 thymus (n = 5 *Morc3*$^{+/+}$, *n* = 16 *Morc3*$^{+/MD41}$ , *n* = 6 *Morc3*$^{MD41/MD41}$, 3 independent experiments). Data are presented as mean with dot, square or triangle as individual values; error bar represents standard deviation (SD). Statistical significance was determined using one-way ANOVA for multiple comparisons, * p≤0.05, **p≤0.01, ***p≤0.001, ****p≤0.0001.

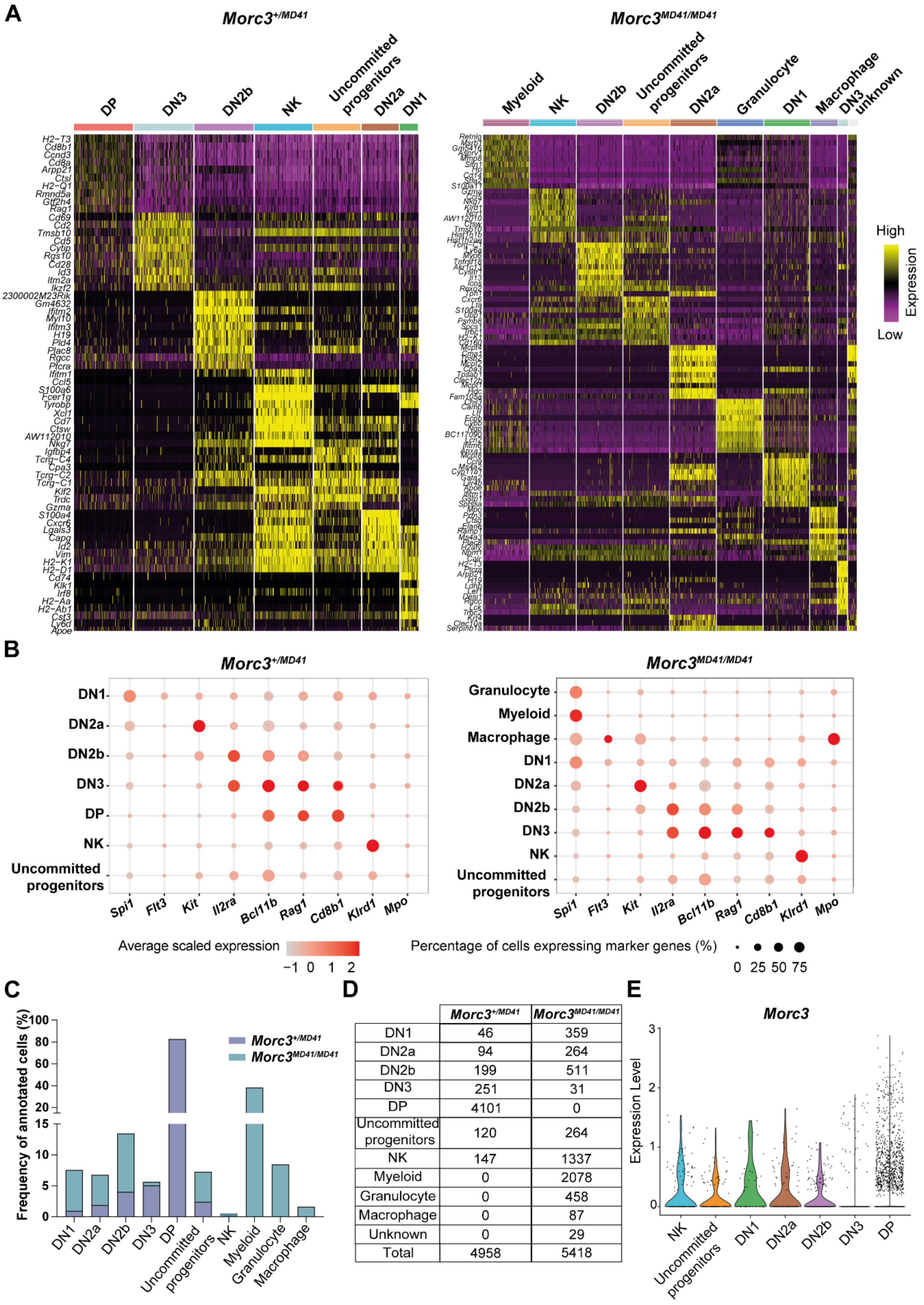


| | *Morc3*+/MD41 | *Morc3*MD41/MD41 |
|---|---|---|
| DN1 | 46 | 359 |
| DN2a | 94 | 264 |
| DN2b | 199 | 511 |
| DN3 | 251 | 31 |
| DP | 4101 | 0 |
| Uncommitted progenitors | 120 | 264 |
| NK | 147 | 1337 |
| Myeloid | 0 | 2078 |
| Granulocyte | 0 | 458 |
| Macrophage | 0 | 87 |
| Unknown | 0 | 29 |
| Total | 4958 | 5418 |

**Supplementary Figure S3. Single-cell RNA-seq – annotation of cell clusters.**

(A) Heatmaps of top 10 marker gene expression in each cluster for each dataset.

(B) Dot plot showing normalized expression of marker genes used to annotate the cell clusters in *Morc3*$^{+/MD41}$ and *Morc3*$^{MD41/MD41}$ E18.5 thymus datasets.

(C) Stacked bar plots showing percentage of annotated cell clusters in *Morc3*$^{+/MD41}$ and *Morc3*$^{MD41/MD41}$ E18.5 thymus datasets.

(D) Table showing total cell number of annotated cell clusters in *Morc3*$^{+/MD41}$ and *Morc3*$^{MD41/MD41}$ E18.5 thymus datasets.

(E) Violin plots showing *Morc3* expression in annotated cell clusters of the *Morc3*$^{+/MD41}$ dataset. Each dot represents a single cell.

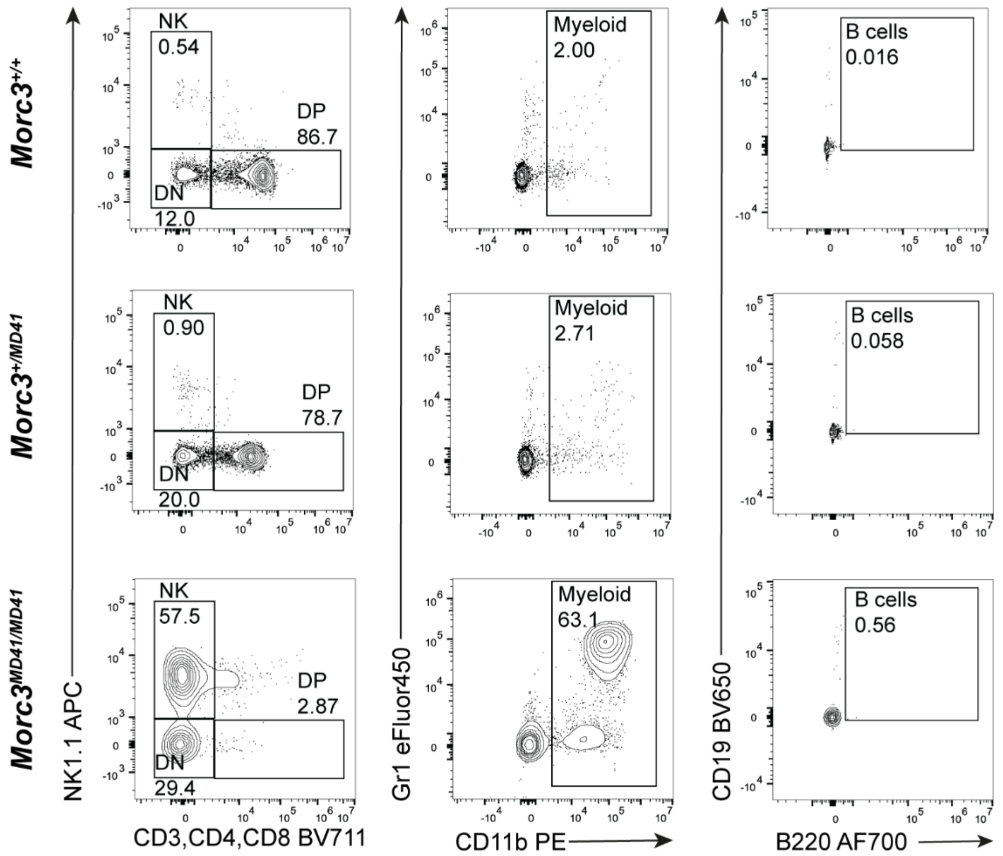


**Supplementary Figure S4. Flowcytometry analysis of lineages in E18.5 thymus.**

Representative flowcytometry plots of DN, DP, NK, myeloid and B cells in *Morc3*$^{+/+}$ , *Morc3*$^{+/MD41}$ and *Morc3*$^{MD41/MD41}$ E18.5 thymus.

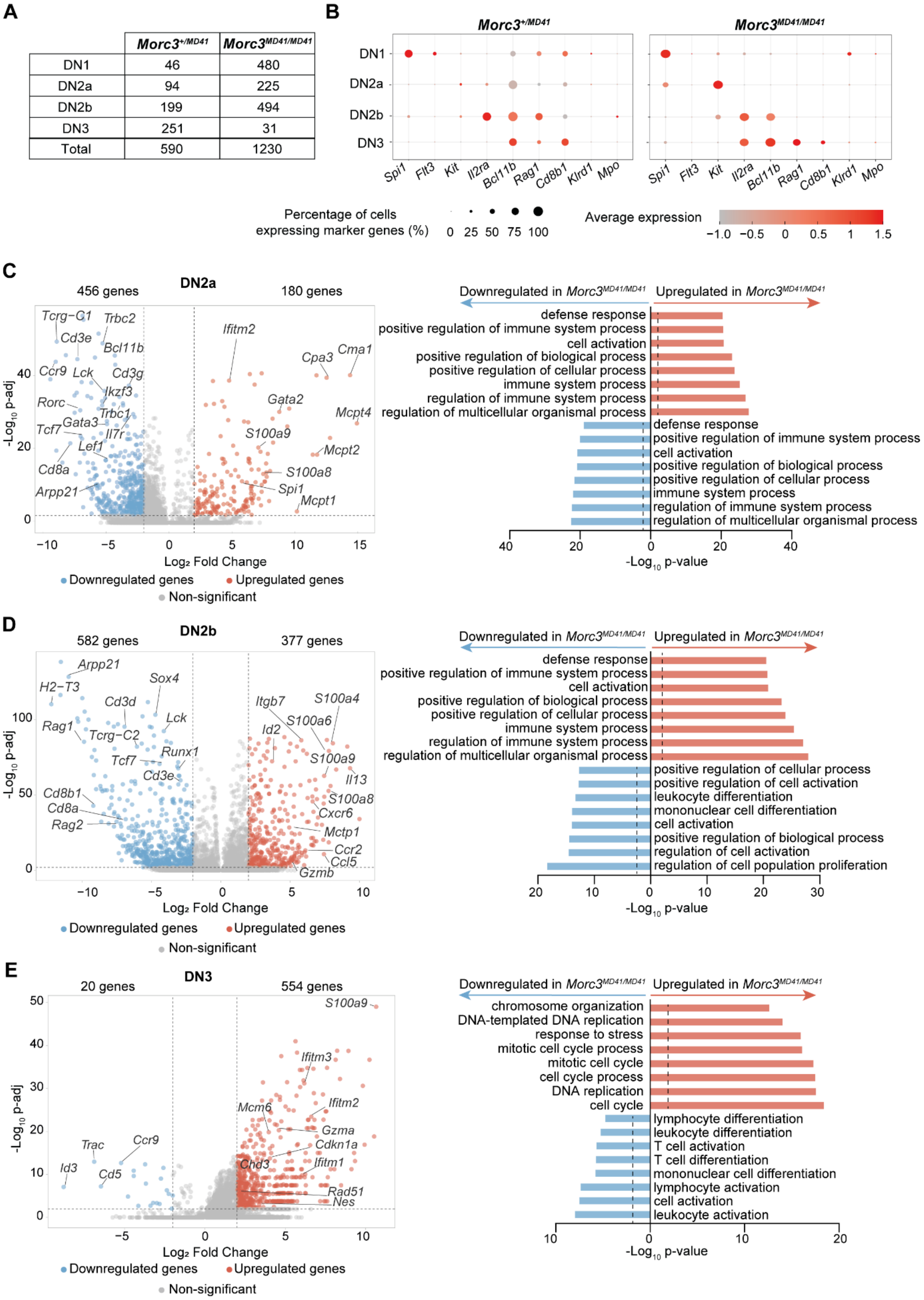


**Supplementary figure S5. Loss of MORC3 affects transcriptome of early T cells.**

(A) Table showing total cell number of annotated cell clusters in *Morc3*$^{+/MD41}$ and *Morc3*$^{MD41/MD41}$ DN subsets.

(B) Dot plot showing normalized expression of marker genes in *Morc3*$^{+/MD41}$ and *Morc3*$^{MD41/MD41}$ subsets of DN cells.

(C) (left) Volcano plot showing differentially expressed genes between *Morc3*$^{+/MD41}$ and *Morc3*$^{MD41/MD41}$ DN2a cells. Genes with |log2 fold change|> 2 and p-adj < 0.01 were considered significant. Red dots indicate upregulated genes in *Morc3*$^{MD41/MD41}$ DN2a. Blue dots indicate downregulated genes in *Morc3*$^{MD41/MD41}$ DN2a cells. Values on top of the plot denote number of upregulated and downregulated genes. The y-axis shows -log10 p-adj and the x-axis log2 fold change. Horizontal dashed line indicates -log p-value of 0,01 (=2), vertical dashed lines indicate log fold change of ±2. (right) Bar graph depicting Gene Ontology enrichment of biological processes in significantly differential gene sets (|log2 fold change|> 2 and p-adj < 0.01) in DN2a cell cluster. Dashed lines indicate the -log p-value of 0,01 (=2).

(D) (left) Volcano plot showing differentially expressed genes between *Morc3*$^{+/MD41}$ and *Morc3*$^{MD41/MD41}$ DN2b cells. Genes with |log2 fold change|> 2 and p-adj < 0.01 were considered significant. Red dots indicate upregulated genes in *Morc3*$^{MD41/MD41}$ DN2b. Blue dots indicate downregulated genes in *Morc3*$^{MD41/MD41}$ DN2b cells. Values in the plot denote number of upregulated and downregulated genes. The y-axis shows -log10 p-adj and the x-axis log2 fold change. Horizontal dashed line indicates -log p-value of 0,01 (=2), vertical dashed lines indicate log fold change of ±2. (right) Bar graph depicting Gene Ontology enrichment of biological processes in significantly differential gene sets (|log2 fold change|> 2 and p-adj < 0.01) in DN2b cell cluster. Dashed lines indicate the -log p-value of 0,01 (=2).

(E) (left) Volcano plot showing differentially expressed genes between *Morc3*$^{+/MD41}$ and *Morc3*$^{MD41/MD41}$ DN3 cells. Genes with |log2 fold change|> 2 and p-adj < 0.01 were considered significant. Red dots indicate upregulated genes in *Morc3*$^{MD41/MD41}$ DN3 cells. Blue dots indicate downregulated genes in *Morc3*$^{MD41/MD41}$ DN3 cells. Values in the plot denote number of upregulated and downregulated genes. The y-axis shows -log10 p-adj and the x-axis log2 fold change. Horizontal dashed line indicates -log p-value of 0,01 (=2), vertical dashed lines indicate log fold change of ±2. (right) Bar graph depicting Gene Ontology enrichment of biological processes in significantly differential gene sets (|log2 fold change|> 2 and p-adj < 0.01) in DN3 cell cluster. Dashed lines indicate the -log p-value of 0,01 (=2).

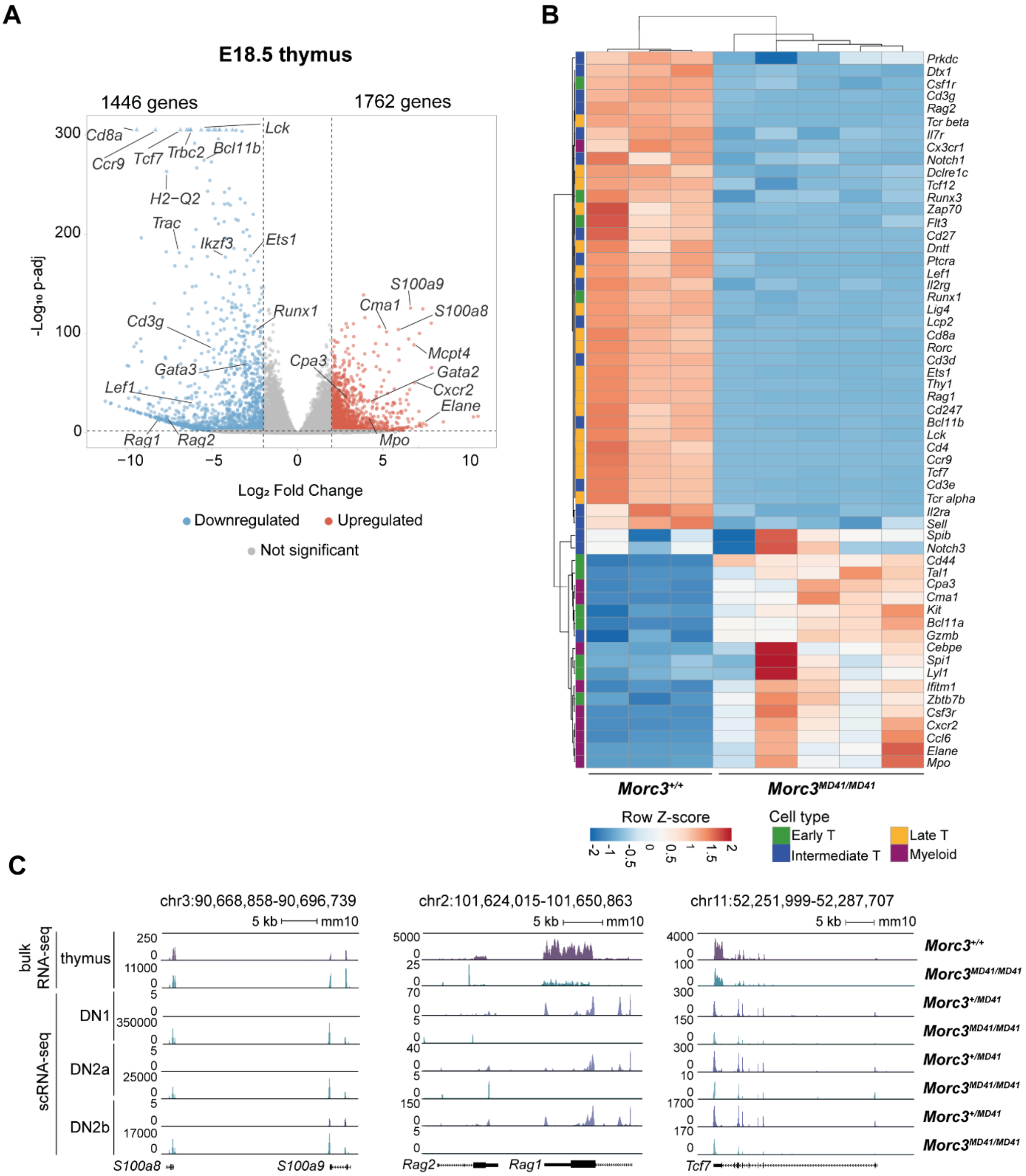


**Supplementary Figure S6. Dysregulated transcriptome in *Morc3$^{MD41/MD41}$* thymus.**

(A) Volcano plot showing differentially expressed genes between *Morc3$^{+/+}$* and *Morc3$^{MD41/MD41}$* E18.5 thymus. (n=3 biological replicates of *Morc3$^{+/+}$* ; n=5 biological replicates of *Morc3$^{MD41/MD41}$* ). Genes with |log2 fold change|> 2 and p-adj < 0.01 were considered significant. Numbers indicate number of upregulated and downregulated genes. Red dots indicate upregulated and blue dots indicate downregulated genes in *Morc3$^{MD41/MD41}$* thymocytes. The y-axis shows -log10 p-adj and the x-axis log2 fold change. Horizontal dashed line indicates -log p-value of 0,01 (=2), vertical dashed lines indicate log fold change of ±2. Triangles indicate out-of-the-range values.

(B) Heatmap depicting expression of marker genes of T cell development (early, intermediate, and late), previously described in [22], and myeloid genes curated from the literature [23]. N=3 biological replicates of *Morc3*$^{+/+}$ ; n = 5 biological replicates of *Morc3*$^{MD41/MD41}$. Colored squares on the left indicate the cell type where the marker gene is mainly expressed.

(C) Genome browser screenshots of *s100a8 and S100a9 loci* (left panel)*, Rag1* and *Rag2* loci (middle panel) and *Tcf7* locus (right panel). Bulk RNA-seq reads in E18.5 thymus and scRNA-seq reads in DN1, DN2a, DN2b cell populations are shown. Scales are adapted for genotyping and cell types.

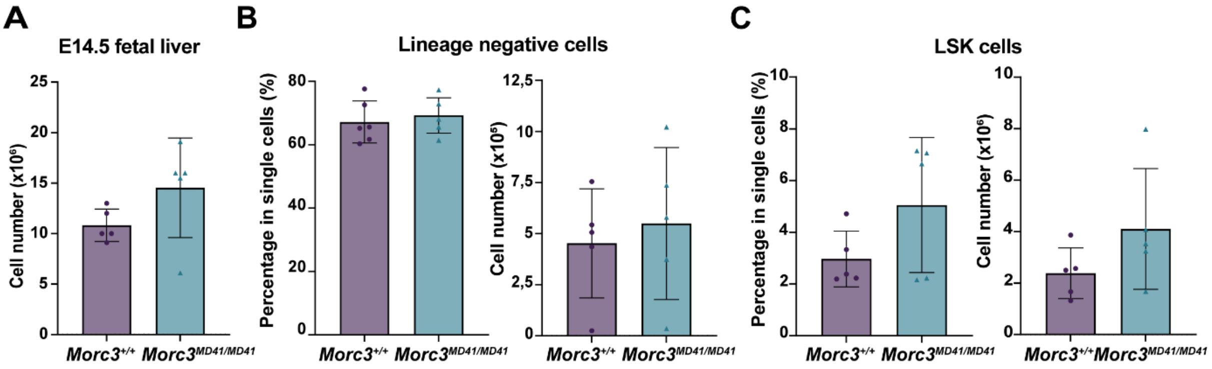


**Supplementary Figure S7. Fetal liver cell analysis**

(A) Bar plot showing the total cell number of E14.5 fetal liver from *Morc3*$^{+/+}$ and *Morc3*$^{MD41/MD41}$ littermates (3 litters, n = 5 *Morc3*$^{+/+}$ and n = 5 *Morc3*$^{MD41/MD41}$ , 3 independent experiments), data are presented as mean with dots/triangles as individual values; error bar represents SD.

(B) (left) Bar plot depicting the percentage within single cells and (right) total cell number of lineage negative cells in E14.5 fetal livers after lineage positive depletion (n = 5 biological replicates per genotype, 3 independent experiments)¸ data are presented as mean with dots/triangles as individual values; error bar represents SD.

(C) (left) Bar plot depicting the percentage within single cells and (right) cell number of LSK (Lineage-, Sca1+, ckit+) cells after lineage depletion (n = 5 biological replicates per genotype, 3 independent experiments)¸ data are presented as mean with dots/triangles as individual values; error bar represents SD.

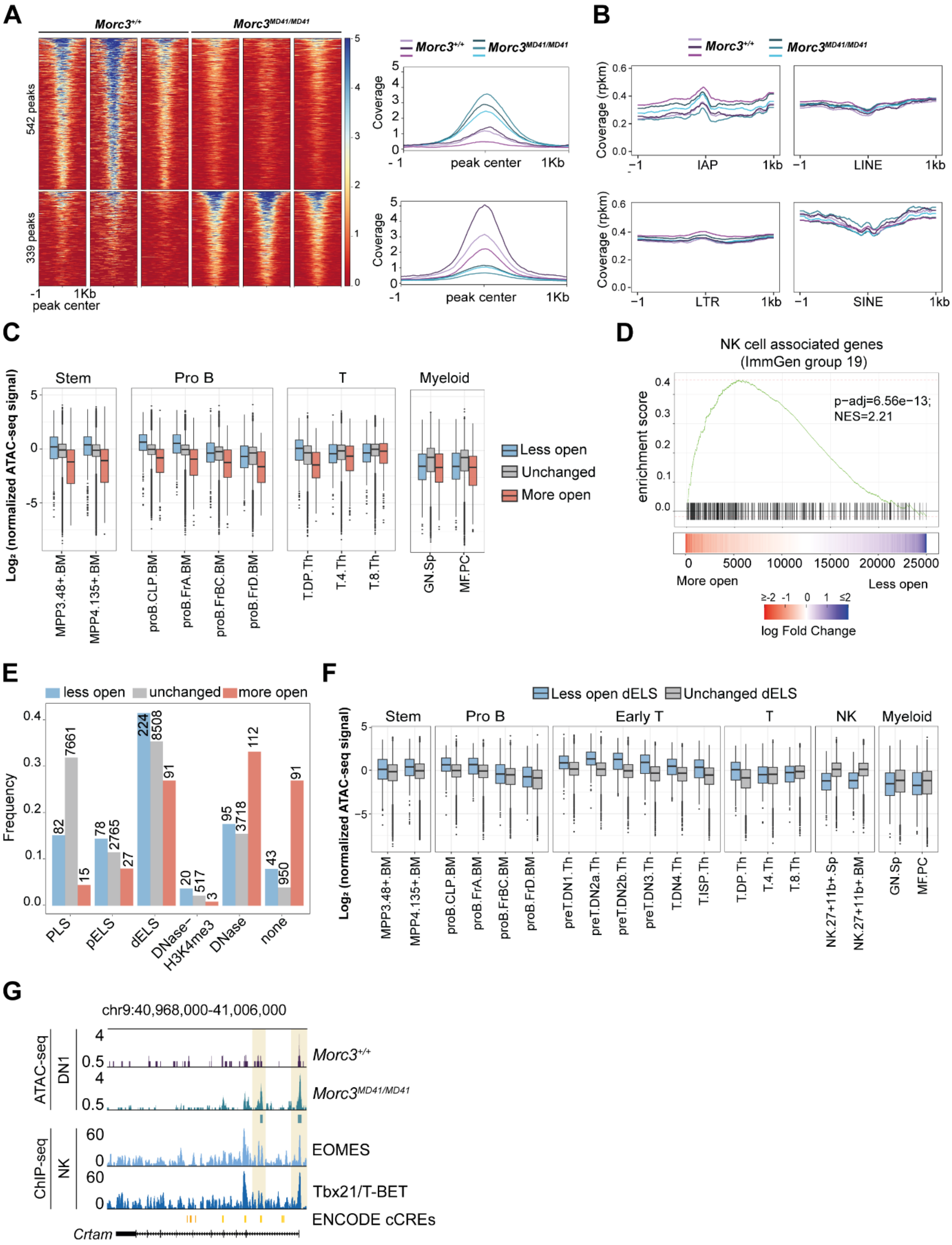


**Supplementary Figure S8. Chromatin accessibility analysis on *Morc3*$^{+/+}$ and *Morc3*$^{MD41/MD41}$ DN1.**

A) (left) Heatmaps depicting ATAC-seq signal of differential accessible peaks (Limma, p-adj < 0.05). (right) Profile plots showing ATAC-seq coverage per replicate over significantly more accessible and less accessible peaks. Biological replicates are shown in a different shade of the same color. Both heatmap and profile plots show ATAC-seq signal over the peak center and 1kb flanking regions (10bp bins and smoothing).

B) Profile plots showing ATAC-seq coverage per replicate over center of indicated transposable elements and 1kb up and downstream (10bp bins and smoothing). All repeatmasker annotations of IAPs were used. For LINE, LTR and SINE elements, 10% highest repeatmasker scores were selected to generate a representative subset. Biological replicates are shown in a different shade of the same color.

C) Box plots depicting normalized ImmGen ATAC-seq signal in less open, unchanged and more open peaks across different immune cell types. The ImmGen ATAC-seq signal was normalized by the mean of ATAC-seq signal over all 90 cell types. Boxes are drawn between 25$^{th}$ and 75$^{th}$ percentiles and the middle line represents the median. Whiskers extend 1.5 * IQR from the end of the box. Points outside this range are plotted individually.

D) GSEA plot of group of peaks associated to genes annotated in ImmGen coarse cluster 19 (annotated as NK cell associated genes). Signed p-values was used to rank peaks. Peaks were associated to genes by taking the closest TSS. Rank 1 was assigned to the most significant more accessible peak, rank 25000 was assigned to the most significant less accessible peak.

E) Bar graph showing frequency and number of less open, unchanged and more open peaks with different annotation derived from the ENCODE data. The annotations are as follows: Promoter Like Signatures (PLS), proximal Enhancer Like Signatures (pELS), distal Enhancer Like Signatures (dELS), regions with both DNase and H3K4me3 (DNase-H3K4me3), only DNase (DNase) and not overlapping with any previous signature (none).

F) Box plots depicting normalized ImmGen ATAC-seq signal in less open, unchanged and more open peaks over ENCODE-annotated dELS across different immune cell types. The ImmGen ATAC-seq signal was normalized by the mean of ATAC-seq signal over all 90 cell types. Boxes are drawn between 25$^{th}$ and 75$^{th}$ percentiles and the middle line represents the median. Whiskers extend 1.5 * IQR from the end of the box. Points outside this range are plotted individually.

G) UCSC genome browser screenshot of the *Crtam* locus. Representative tracks for ATAC-seq from DN1 cells, published ChIP-seq (EOMES: GSM5134535; Tbx21/T-BET: GSM5134533) and ENCODE cis-regulatory elements (CREs) in orange boxes are shown. Yellow shading indicates representative differential ATAC-seq peaks.

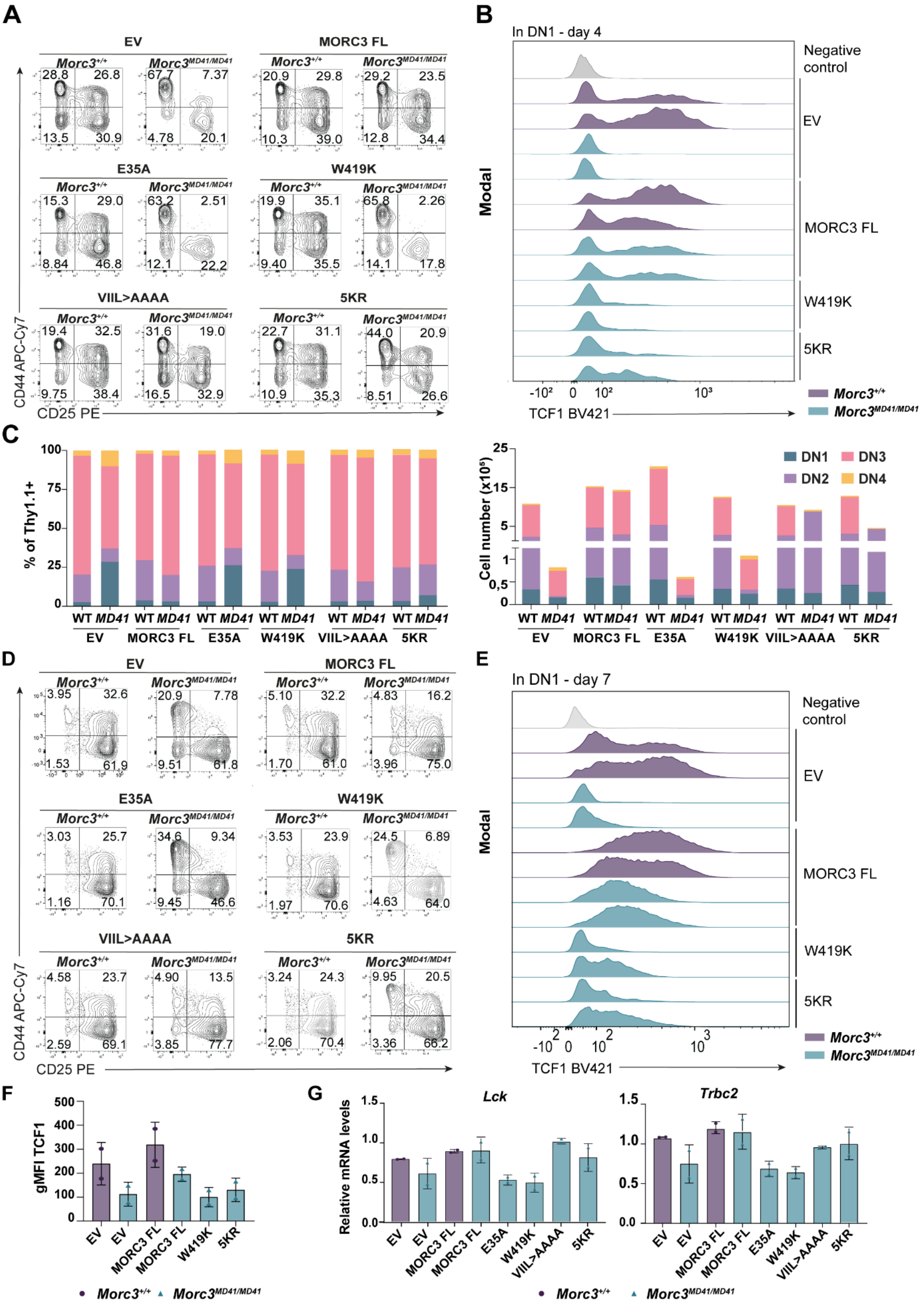

**Supplementary Figure S9. MORC3 ATP hydrolysis and H3K4me binder mutant proteins fail to fully rescue T cell development defect in *Morc3*$^{MD41/MD41}$ cells.**

(A) Representative flowcytometry plots of DN1, DN2, DN3 and DN4 in OP9-DL1 co-cultures of *Morc3*$^{+/+}$ and *Morc3*$^{MD41/MD41}$ cells transduced with empty vector (EV), MORC3 FL, MORC3 mutants (E35A, W419K, VIIL>AAAA, 5KR), on day 4. All cells are pre-gated on SSC-A/FSC-A, singles, live cells (7AAD-), Lin-, CD45.1+, GFP+ (only for EV transduced cells), Thy1.1+.

(B) Histogram depicting TCF1 intracellular expression in DN1 cells measured by flowcytometry after 4 days of OP9-DL1 co-culture of *Morc3*$^{+/+}$ and *Morc3*$^{MD41/MD41}$ cells transduced with EV, MORC3 FL, W419K or 5KR Morc3 mutants. The negative control is *Morc3*$^{+/+}$ CD45.1- cells. All cells are pre-gated on SSC-A/FSC-A, singles, live cells (7AAD-), Lin-, CD45.1+, GFP+ (only for EV transduced cells), Thy1.1+, CD44+CD25-.

(C) Stacked bar chart indicating the percentage (left) and total cell count (right) of DN1, DN2, DN3 and DN4 cells of *Morc3*$^{+/+}$ and *Morc3*$^{MD41/MD41}$ cells transduced with EV, MORC3 FL, MORC3 mutants (E35A, W419K, VIIL>AAAA, 5KR), on day 7 after OP9-DL1 co-culture system. (n = 8 GFP control, MORC3 FL; n = 6 W419K, 5KR; n = 4 VIIL>AAA, E35A, in 4 independent experiments). All cells are pre-gated on SSC-A/FSC-A, singles, live cells (7AAD-), Lin-, CD45.1+, GFP+ (only for EV transduced cells), Thy1.1+.

(D) Representative flowcytometry plots of DN1, DN2, DN3 and DN4 in OP9-DL1 co-cultures of *Morc3*$^{+/+}$ and *Morc3*$^{MD41/MD41}$ cells transduced with EV, MORC3 FL and MORC3 mutants (E35A, W419K, VIIL>AAAA, 5KR), on day 7.

(E) Histogram depicting TCF1 intracellular expression in DN1 cells measured by flowcytometry after 7 days of OP9-DL1 co-culture of *Morc3*$^{+/+}$ and *Morc3*$^{MD41/MD41}$ cells transduced with EV, MORC3 FL, W419K or 5KR MORC3 mutants. The negative control is *Morc3*$^{+/+}$ CD45.1- cells. (n=2 biological replicates per genotype). All cells are pre-gated on SSC-A/FSC-A, singles, live cells (7AAD-), Lin-, CD45.1+, GFP+ (only for EV transduced cells), Thy1.1+, CD44+CD25-.

(F) Bar plot showing geometric mean fluorescent intensity (gMFI) of TCF1 expression in DN1 cells after 7 days of co-culture of *Morc3*$^{+/+}$ and *Morc3*$^{MD41/MD41}$ cells transduced with EV, Morc3 FL, W419K or 5KR MORC3 mutants. gMFI was calculated by subtracting gMFI of TCF1 expression in negative control. n = 2 biological replicates per genotype. Data are presented as mean with dot/triangle as individual values; error bar represents SD.

(G) RT-qPCR analysis of relative *Lck* and *Trbc2* mRNA levels in *Morc3*$^{+/+}$ and *Morc3*$^{MD41/MD41}$ cells after 7 days of OP9-DL1 co-culture. Data are normalized to *Ptprc*; n = 2 biological replicates per genotype; data are presented as mean with dot/triangle as individual values; error bar represents SD.

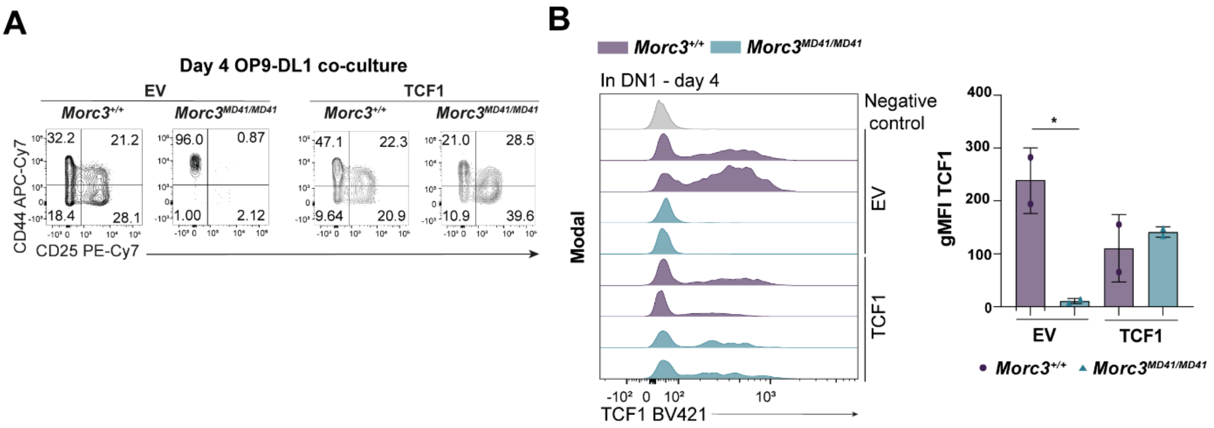


**Supplementary Figure S10. Ectopic TCF1 expression in *Morc3$^{MD41/MD41}$* cells.**

(A) Representative flowcytometry plots of DN1, DN2, DN3 and DN4 in *Morc3$^{+/+}$* and *Morc3$^{MD41/MD41}$* cells transduced with empty vector (EV) or TCF1 after 4 days of OP9-DL1 co-culture. All cells are pre-gated on SSC-A/FSC-A, singles, Lin-, CD45.1+, GFP+, Thy1.1+.

(B) (left) Histogram depicting TCF1 intracellular expression in DN1 cells (gated on SSC-A/FSC-A, singles, Lin-, CD45.1+, Thy1.1+, CD44+CD25-) measured by flowcytometry after 4 days of OP9-DL1 co-culture of *Morc3$^{+/+}$* and *Morc3$^{MD41/MD41}$* cells transduced with EV or TCF1 virus. The negative control is *Morc3$^{+/+}$* CD45.1- cells. (right) Bar plot showing geometric mean fluorescent intensity (gMFI) of TCF1 expression in DN1 cells as depicted in the left panel. The gMFI was calculated by subtracting gMFI of TCF1 expression in negative control. n = 2 biological replicates per genotype. Data are presented as mean with dot/triangle as individual values; error bar represents SD.

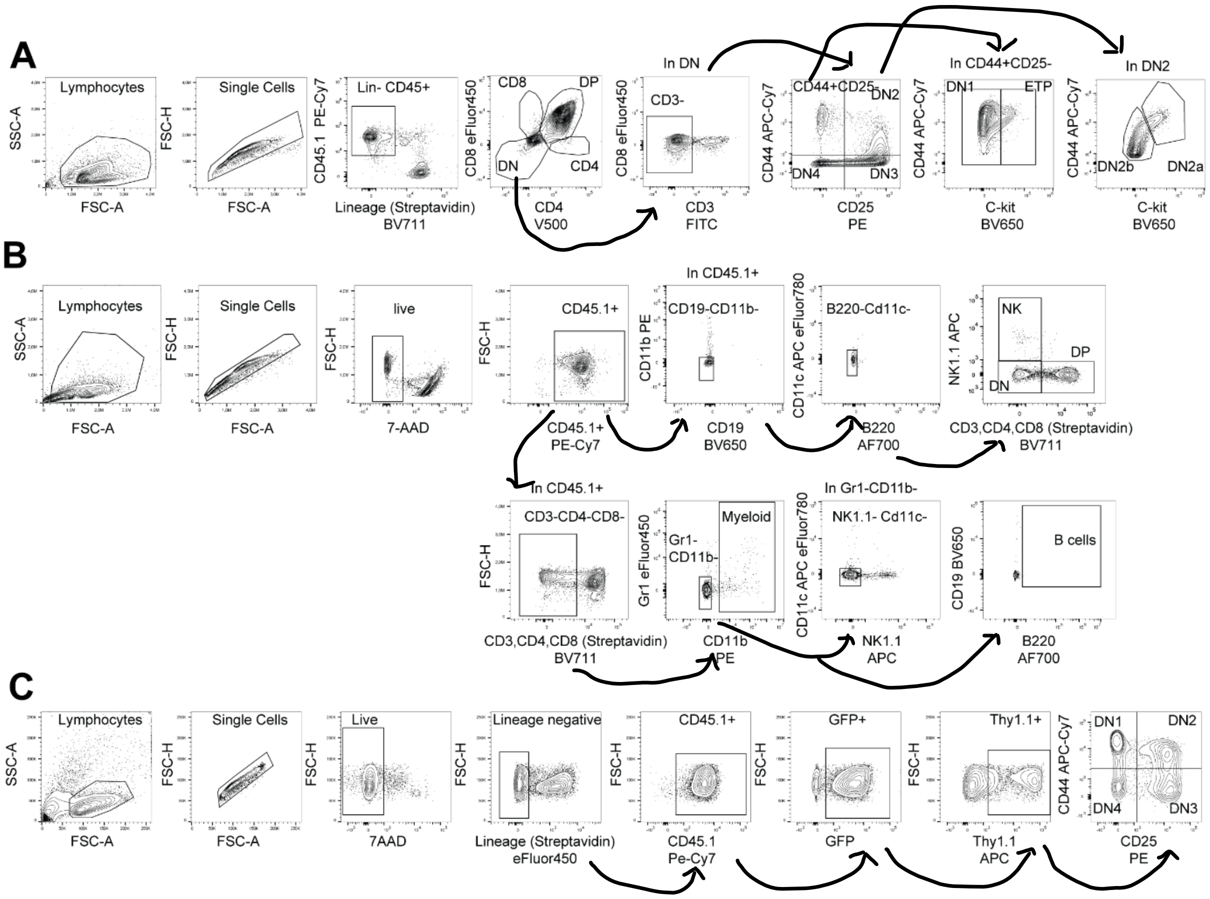


**Supplementary Figure S11. Gating strategy of flowcytometry analysis.**

(A) Gating strategy for T cell populations in E18.5 thymus.

(B) Gating strategy for lineages in E18.5 thymus.

(C) Gating strategy for DN cells in OP9-DL1 co-culture system (GFP gate included only in cells transduced with EV retrovirus).
